# Differentiation of cortical brain organoids and optic nerve-like structures from retinal confluent cultures of pluripotent stem cells

**DOI:** 10.1101/2021.05.16.444356

**Authors:** Milan Fernando, Scott Lee, Jesse R. Wark, Di Xiao, Hani J. Kim, Grady C. Smith, Ted Wong, Erdahl T. Teber, Robin R. Ali, Pengyi Yang, Mark E. Graham, Anai Gonzalez-Cordero

**Author notes:** **Corresponding author:** Dr Anai Gonzalez Cordero, Stem Cell Medicine Group, Children’s Medical Research Institute and University of Sydney, 214 Hawkesbury Road, Westmead, 2145, NSW, Australia. these two authors contributed equally to the work (Milan Fernando, Scott Lee).

## Abstract

Advances in the study of neurological conditions have been possible due to induced pluripotent stem cell technologies and the generation of neural cell types and organoids. Numerous studies have described the generation of neural ectoderm-derived retinal and brain structures from pluripotent stem cells. However, the field is still troubled by technical challenges, including high culture costs and organoid-to-organoid variability. Here, we describe a simple and economical protocol that reproducibly gives rise to the neural retina and cortical brain regions from confluent cultures of stem cells. The spontaneously generated cortical organoids were isolated and cultured in suspension conditions for maturation and are transcriptionally comparable to organoids generated by other methods and to human foetal cortex. Furthermore, these organoids show spontaneous functional network activity with proteomic analysis and electron microscopy demonstrating the presence of synaptic components and maturity. The generation of retinal and brain organoids in close proximity also enabled their mutual isolation. Further culture of this complex organoid system demonstrated the formation of optic nerve-like structures connecting retinal and brain organoids, which might facilitate the investigation of the mechanisms of neurological diseases of the eye and brain.

## Introduction

The rapidly progressing field of human pluripotent stem cells (hPSCs), including embryonic (ES) and induced pluripotent stem (iPS) cells, and their derivative organoids continue to provide new insights into basic biology, human development, modelling of human diseases and discovery of innovative treatments. Neural differentiation, in particular, has been extensively studied, improving our understanding of the mechanism of neurodevelopmental conditions^1,2,3^. Large numbers of neurons and astrocytes are able to be generated using two-dimensional (2D) cultures derived from hPSCs^4,5^ and these classical differentiation cultures were later optimised to three-dimensional (3D) suspension methods that better recapitulate the physiological niche and environment of the developing human brain^6^.

In the developing central nervous system (CNS), the eye and the brain form as an extension of the forebrain diencephalic and telencephalic region, respectively^7^. Developmental biology studies have helped infer the molecular basis of this patterning process and facilitated the establishment of numerous differentiation protocols, which generate miniaturised versions of 3D organoids from PSCs. Brain organoids replicate specific brain regions or whole cerebral areas, with both occasionally developing eye regions^8 9,10^.

Differentiation protocols are usually classified as either guided/direct or non-guided/undirected cultures, based on the necessity for guided differentiation using growth factors or their absence in non-guided spontaneous differentiation, which relies on endogenous self-forming ability of the cells^11^. Brain organoids generated using these varied methodologies have been comprehensively characterised using transcriptome analysis by RNA sequencing. Single cell RNA sequencing (scRNA-seq) has elucidated the cellular composition of organoids and reproducibility of organoid protocols^12,13,1,10,14-16^. However, the differentiation of PSC-derived brain organoids presents challenges, such as variability within and between organoid batches and high culture costs. This lack of reproducibility demonstrates the necessity of novel differentiation approaches. Furthermore, unbiased omics, including proteome studies, together with robust measurements of neuronal activity, are critical to establish organoid variability, maturation and functionality.

Diseases of the eye and the brain are now understood to be more intertwined than previously thought. Studies of common conditions, such as Alzheimer’s disease and glaucoma have demonstrated neurodegenerative changes and disease traits in both brain and eye regions^17,18^. Complex organoids have the potential to provide useful models of these human disease *in vitro* with the proviso that they faithfully recapitulate retinal development, morphology and maturation. Therefore, improved formation of hPSC-derived retinal-brain connection through an optic nerve is essential for effective degenerative disease modelling.

Here we hypothesised that it is possible to reproducibly generate functional brain organoids from retinal confluent cultures of PSCs. We also asked the question if the retinal and brain organoids developing in suspension culture together would form a complex retina-brain organoid system that would enable the formation of an optic nerve-like structure.

We demonstrated here that the non-guided, simple and economical differentiation protocol, previously described to spontaneously generate retinal vesicles from a confluent culture of hPSCs^19-21^, also generated brain organoids alongside the differentiation of retinal vesicles. The ease of precisely locating cortical organoids, due to morphology and their proximity to retinal vesicles, reduced organoid variability. Brain organoids were characterised as dorsal cortical organoids which, when further cultured in 3D suspension, matured into functional organoids. A systematic comparison of scRNA-seq datasets revealed a close similarity of our organoids with other dorsal patterned hPSC-derived brain organoids^16^ and proteomic analysis of late-stage organoids revealed the presence of numerous synaptic markers. Organoids electrophysiological activity was dependent on organoids being cultured in relevant basal medium^22^ with proteomics providing insights into why this physiological environment aids neurophysiological activity. Finally, the generation of retinal and cortical organoids facilitated the isolation of both structures for 3D culturing, forming a complex organoid system. Notably, when cultured in suspension, these retinal-brain organoids maintained the natural association created in the dish during their spontaneous development which enabled the formation of optic nerve-like structures between the two organoids.

## Results

### Generation of self-forming retinal and dorsal cortical brain organoids from Confluent 2D/3D cultures

The retina is an extension of the CNS arising from the forebrain region in the developing embryo (**Figure 1a)**. Indeed, the differentiation of whole brain cerebral organoids containing retinal structures has been demonstrated^8,10^. **Figure 1b** schematic shows that proneural induction of confluent cultures of PSCs spontaneously generate pigmented islands of retinal pigment epithelium (RPE) from which retinal vesicles appear ^19,21^. Further analysis of other structures forming in this 2D/3D environment highlighted the presence of 3D regions containing clear neuronal rosettes forming in close proximity to retinal vesicles. Based on the neuronal culture environment, and their morphological characteristics, we hypothesised that these were brain vesicles. To fully characterise these structures, we manually excised these regions to be grown in 3D suspension for maturation in culture, as previously described for their retinal counterparts ^21^.

**Figure 1.**
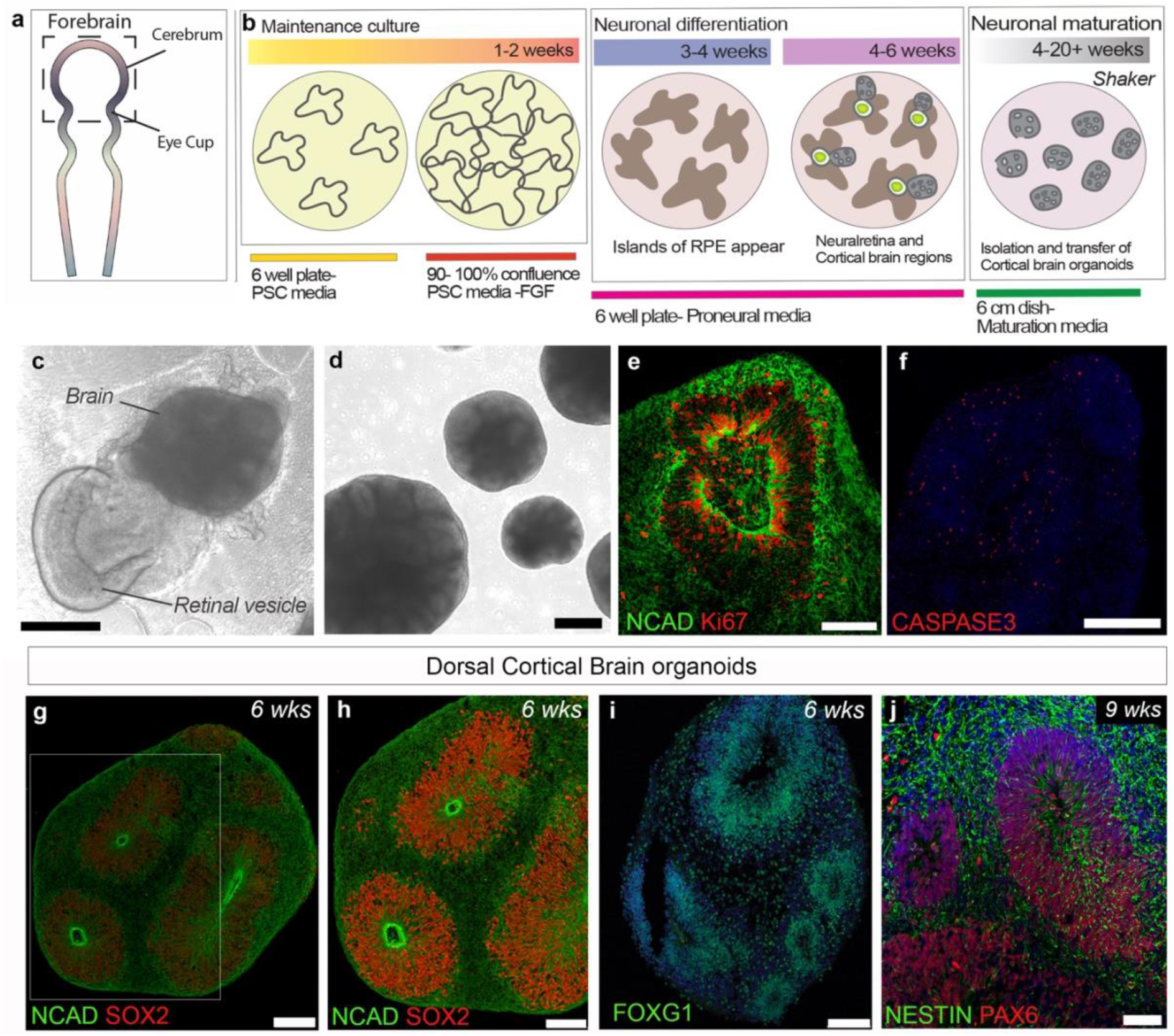
Confluent PSC differentiation protocol gives rise to cortical and retinal organoids. **a**. Schematic illustrating neural tube formation and the formation of both cerebellum and eye cups from forebrain. **b**. Schematic summarising the 2D/3D differentiation protocol timeline for both retinal and cortical organoids formation as well as media composition and maturation in suspension. **c**. Representative image of a retinal vesicle and brain organoid in 2D culture. **d**. Bright-field image showing floating brain organoids, with typical neural rosettes, following manually excision from 2D culture. **e-j**. Immunohistochemistry images of 6 -9 weeks old brain organoids. **e, f**. Brain organoids showing Ki67 proliferative NCAD positive neuroepithelium and CASPASE 3 apoptotic cells. **g-j**. Cortical origin of brain organoids is confirmed by the presence of SOX2, FOXG1 and PAX6 positive neural precursors markers. Scale bars, 50 μm (j), 75 μm (h), 100 μm (e, g, i), 250 μm (f).

Following a neural induction period of 4-6 weeks in culture, neuroretinal vesicles and neural rosette structures appeared (**Figure 1c**). These areas were manually dissected and grown in suspension in retinal media from 7 weeks onwards to allow for maturation in a media composition previously described in Lancaster et al., 2013 (**Figure 1d**). By 6 weeks of differentiation, 39% of all organoids observed in cultures were retinal vesicles while 60% were neuronal organoids (n=10 2D/3D differentiation batches) demonstrating that a proportion of cortical organoids are generated independently and are not in proximity to retinal organoids. Immunohistochemistry (IHC) of 6-week old organoids revealed highly proliferative KI67/NCAD positive neuroepithelium regions (**Figure 1e**). Active CASPASE3 cells were also observed in the organoids. To mitigate this phenotype, organoids were further cultured on previously-described shaking platforms (**Figure 1f**). Well-formed neuroepithelium regions contained numerous SOX2/NCAD positive neural progenitor cells (**Figure 1g, h)** that were also immuno-positive for cortical FOXG1 and NESTIN/PAX6 markers (**Figure 1j, S1a and SMovie 1**). These cortical organoids were negative for ventral cortex specific marker, NKX2.1 (**Figure S1b**), as opposed to whole brain cerebral organoids (**Figure S1c**), confirming the dorsal cortical origin of these organoids.

The presence of RPE and typical retinal vesicles facilitated the separation of dorsal cortical brain organoids under bright-field microscopy from other forebrain-like neuroepithelia which also form in these cultures (**Figure S1d-f**). These dorsal brain organoids could be easily distinguished in the 2D/3D confluent cultures enabling the isolation of a population of dorsal cortical organoids for further maturation and therefore minimising variability of mature cultures. We observed successful differentiation and these culture features across a number of iPS stem cell lines (**Figure S2**).

### Cortical organoids differentiate into Cortical neurons and Glial cell types

Next, IHC confirmed the presence of cortical plate markers TBR1 which co-expressed with CTIP2-positive cells in the cortical organoids (**Figure 2a-c**). By 10 weeks of culture, CTIP2 regions also contained SATB2-positive cells (**Figure 2e, f**). From 8 to 12 weeks in culture, organoids increased in size significantly from 2000μm (± 560) to 2600μm (± 474) in diameter (**Figure 2g**, n=20-25 organoids, N=3 differentiation batches; mean ± SD, p=0.0002, unpaired two-tailed *t* test). This differentiation protocol also supported the differentiation of S100B-(**Figure 2g)** and GFAP-positive astrocytes that were present in similar percentages to TUJ1-positive neurons (**Figure 2h, i**, 75% (±14%) neurons versus 57% (±17%) glial cells; n = 10 images; N = 3-4 differentiation batches; mean ± SD, p <0.0001, unpaired two-tailed *t* test). Further IHC analysis at 15 weeks (∼3.5 months) in culture demonstrated the presence of more mature inhibitory gamma aminobutyric acid (GABAergic) neurons (**Figure 2j, k** (high magnification of boxed area in A)) and their CALRETININ-positive subtypes (**Figure 2l**). Furthermore, 3D imaging of the whole-brain organoid by Lightsheet microscopy confirmed the presence of numerous inhibitory CALRETININ-positive neurons and GFAP-positive astroglial cell types (**SMovie 2**).

**Figure 2.**
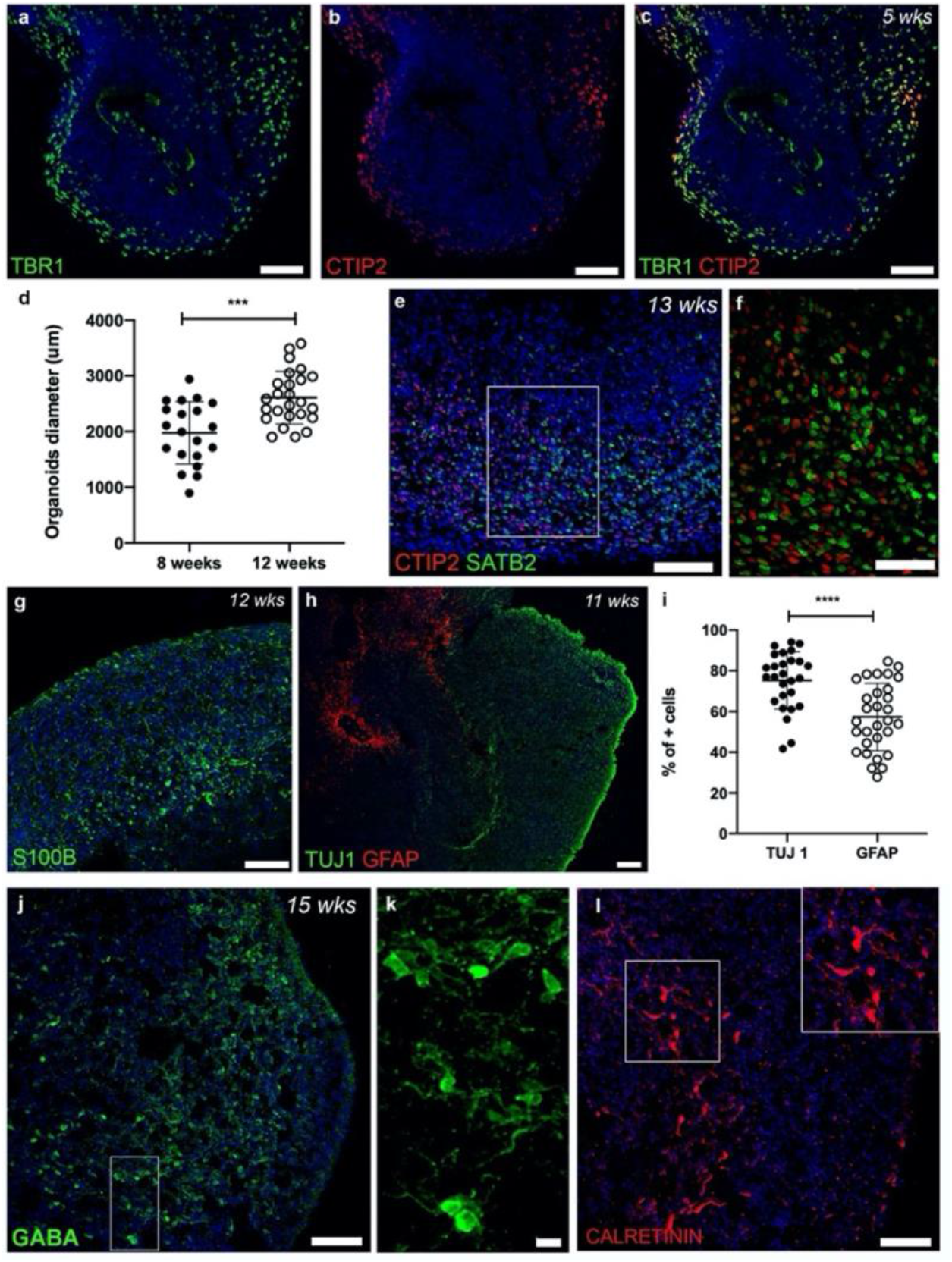
Cortical brain organoids generate cortical plate neurons and glial cells. **a-c**. Representative immunohistochemistry images of week 5 cortical organoids showing the presence of TBR1 and CTIP2 cortical plate neurons. **d**. Graph showing cortical organoids diameter and their significant size increase with days in culture (n= 30 organoids, N= 3 differentiation batches, mean ± SD, unpaired two-tailed *t* test). **e-h**. Representative images of organoids showing CTIP2 and SATB2 positive cells (**e, f**; high magnification image of inset in **e** shown in **f**) and S100B and GFAP glial cells (**g, h**). **i**. Graph showing the percentage of cells expressing TUJ1 and GFAP per area (mm2) in 11 weeks old cells. (n = 10 images from 2-3 organoids, N= 3 differentiation batches, mean ± SD, unpaired two-tailed *t* test). **j-l**. Week 15 cortical organoid showing GABA and CALRETININ positive inhibitory neurons, **k** shows high magnification image of inset in **j**. Scale bars, 15 µm (k), 50 µm (f), 75 µm (a-c), 100 µm (e, g, h, j, l).

### Cortical Organoids from 2D/3D confluent cultures have similar cell type compositions to other brain organoids

Next, we performed scRNA-seq analysis using the 10X genomics platform to further investigate the cell-type composition of these dorsal cortical organoids and to establish their similarity to other published brain organoid scRNA-seq datasets, which were derived from various directed and undirected protocols. We used scClassify^23^, a machine learning-based method, to annotate cell types that were present in our scRNA-seq cortical organoids (**Figure 3a**) and compared the composition of cell types in 3-month-old cortical organoid (CO) we generated against those from 3- and 6-month-old brain organoids and cell types from gestational week 12 (GW12) pre-frontal lobe (PFL) foetal brain (**Figure 3b**).

**Figure 3.**
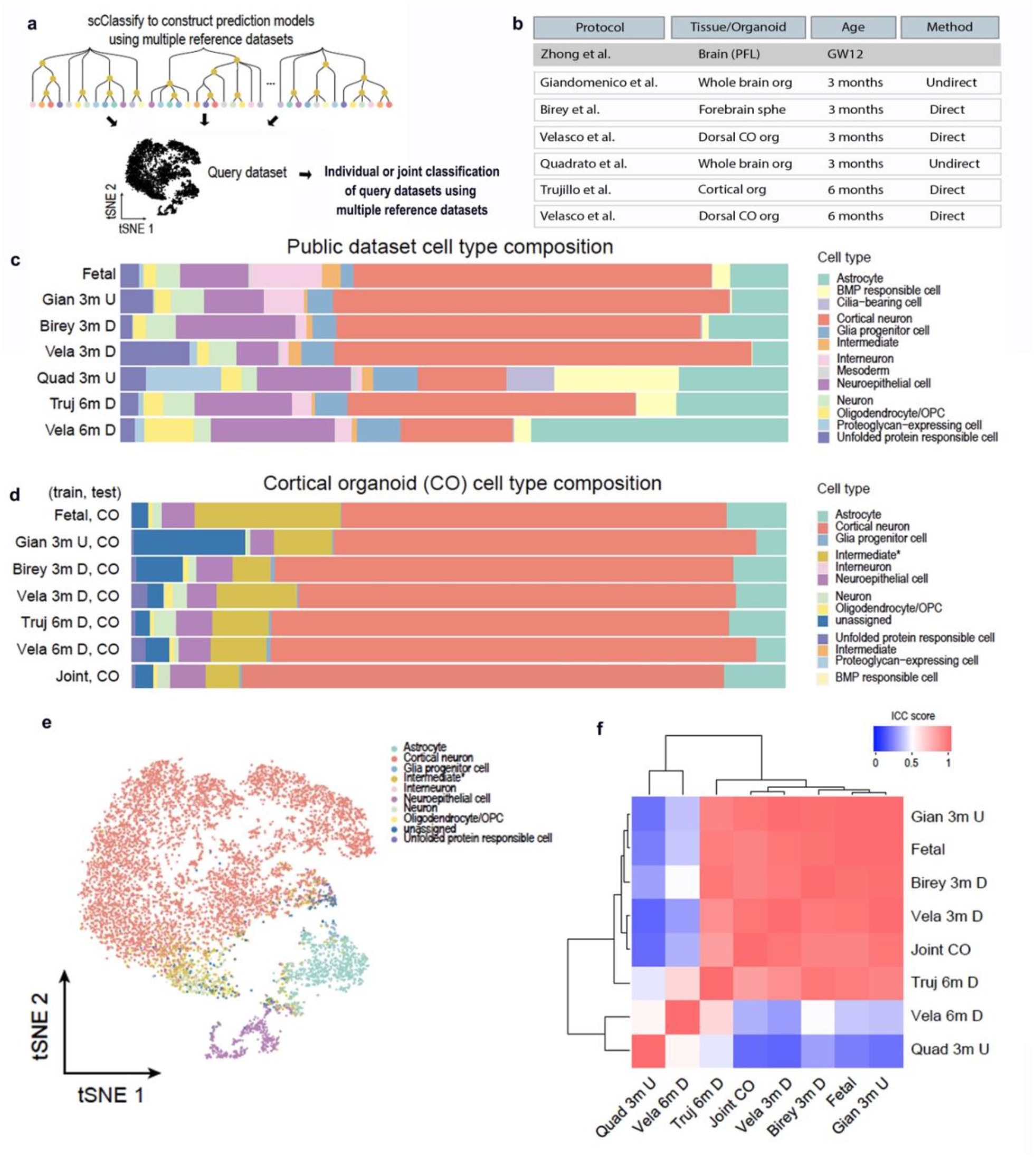
Comparison of cortical organoids and scRNA-seq available dataset. **a**. Schematic illustrating scClassify computational analysis to categorise a dataset using multiple reference datasets. **b**. Table summarising the public scRNA-seq reference datasets included in comparison and their differentiation protocols. **c**. Cell types composition in public datasets based on the annotation from their original studies. **d**. Cell-type composition in cortical organoids (CO) scRNA-seq dataset predicted by scClassify trained by each or all (i.e. Joint) public datasets. **e**. tSNE plot illustrating the cell types composition in CO based on scClassify annotation using Joint training. **f**. Interclass correlation heatmap showing the agreement between cell-type compositions among different organoids and a fetal brain.

First, we compared the composition of cell types by using the cell type labels from the original authors for each of the published datasets, summarised in the table in **Figure 3b**. Although the proportions of each cell type varied across protocols, the overall cell type compositions were similar. This was independent of the protocol, their method of differentiation (directed or undirected) and age of the organoid (**Figure 3c**). In this analysis, Quadrato et al data (Quad 3M U) showed the most diverse cell-type composition.

Next, we utilised scClassify to annotate cells in our CO scRNA-seq dataset by training the classification model using either each individual public dataset as a reference or using all datasets jointly (refer to as joint training) (**Figure 3d**). The predicted cell-type composition in CO scRNA-seq data was plotted with respect to the training data, and, irrespective of the training dataset, a similar cell-type composition was observed in our COs (**Figure 3d**). In agreement with this, the expression profiles of key marker genes for each cell type population were also largely consistent, irrespective of the training dataset (**Figure S3)**. BMP responsive cells, intermediate (bright orange) and proteoglycan-expressing cell types were absent from our dataset. Note that cells are annotated as “unassigned” when they could not be classified to any cell type. Cells are annotated as “intermediate” when they could not be classified to a specific cell type but an intermediate among multiple cell types. This intermediate population (denoted with “*”) is different from the intermediate cell type annotated in the training datasets (bright orange). Since the scClassify jointly trained using multiple reference datasets reduces both unassigned and intermediate* classification, the classification result from the jointly trained model was used in subsequent comparison. Organoid cell types were clustered in a tSNE plot showing that cortical neurons were the most abundant cell type (**Figure 3e**). We then measured the agreement between different cell-type compositions across the multiple datasets using intraclass correlation (ICC) and visualised these results (**Figure 3f**). Three major correlating groups were evident in relation to cell-type compositions: (1) Velasco *et al*., 6 mo., directed (Vela 6m D); (2) Quadrato *et al*., 3 mo., undirected (Quad 3m U) and; (3) the remaining protocols, which included our 2D/3D COs. In accordance with **Figure 3c**, except for Quad 3m U, all 3m organoids and the GW12 foetal sample cluster together, suggesting similar cell-type composition. Notably, our organoids closely resemble the guided differentiation of dorsally patterned forebrain organoids (Vela 3M D), known for their reproducibility^16^. Cell-type proportions were more diverse when Quad 3m U was used as training data, suggesting a unique cell-type composition in this dataset. In fact, Quad 3m U cell-type annotations clustered as a group with Vela 6m D separate from Trujillo *et al*., 6 mo., directed (Truj 6m D) and the remaining 3 mo. organoids. Therefore, whilst cell-type composition of Vela 6m D appeared to be different from those of 3m organoids, the composition of Truj 6m D was more similar to 3m organoids. These results demonstrated the transcriptional similarities between cortical organoids from spontaneous 2D/3D cultures and previous described brain organoids and the foetal nervous system.

### Proteome analysis of brain organoids highlights increased abundance of proteins related to synaptic transmission

The proteomes of iPSCs and cortical organoids were surveyed to a depth of 6,244 and 5,719 proteins, respectively (after filtering and counting only unique genes). There were 4,444 proteins present shared between the iPSC and cortical organoid lists. Gene ontology enrichment analysis was performed with a focus on biological process and KEGG pathway terms (**Figure 4a-d**). **Figure 4a** shows the top thirty biological process terms that had the largest difference in significance between iPSCs and cortical organoids. Terms related to the cell cycle, chromosomes and DNA were more likely to be enriched for iPSCs, whereas terms related to the synapse, synaptic vesicles, vesical transport, axon/dendrite development and neuron projection were more enriched for cortical organoids. Since we are interested in neuronal differentiation in organoids, we extracted the top five synaptic and development terms (not already shown in **Figure 4a**) from the list of significant biological process terms (**Table S2**). For example, the terms “postsynapse organisation” and “regulation of postsynaptic membrane neurotransmitter receptor levels” were enriched for cortical organoids (**Figure 4b**). Neuron development and dendritic spine development were enriched for cortical organoids (**Figure 4c**). The KEGG pathway terms ribosome biogenesis in eukaryotes, DNA replication, cell cycle, and basal transcription factors were more enriched for iPSCs while axon guidance, neurotrophin signalling pathway and long-term potentiation terms were enriched for cortical organoids (**Figure 4d**). Examples of detected proteins that were exclusive to the iPSCs and are involved in stem cell proliferation and maintenance include proteins encoded by *LIN28A, BMPR1A, FGFR1, SOX2, ARID1A* and *RTF1* (**Figure 4e**). Conversely, GABA receptor subunits (encoded by *GABBR1* and *GABRG2*), a glutamate receptor subunit (encoded by *GRIA1*), a mediator of postsynaptic plasticity (encoded by *SYNGAP1*), a trans-synaptic protein (encoded by *NRXN1*) and two synaptic vesicle associated proteins (encoded by *SYN1* and *SYP*) are examples of proteins detected exclusively in cortical organoids (**Figure 4e**). The presence of these proteins and enrichment of relevant gene ontology terms confirms there was strong proteomic evidence that cortical organoids differentiated towards the neuronal- and away from a stem cell-phenotype.

**Figure 4.**
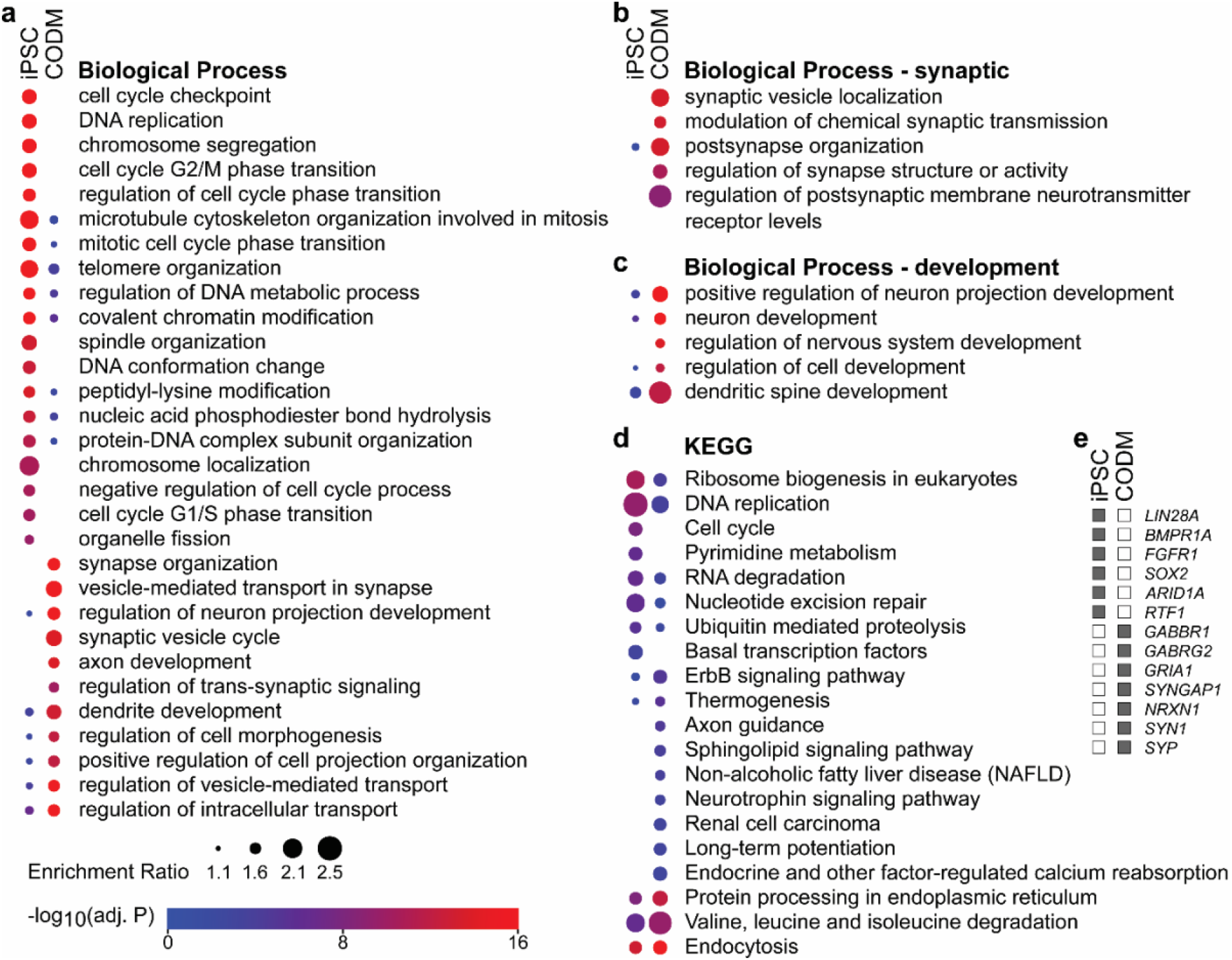
Comparison of the iPSC and cortical organoid proteomes using gene ontology enrichment analysis. **a**. Comparison of biological process terms with the largest difference in probability of enrichment for the genes encoding proteins detected in the iPSC and cortical organoid proteomes, the latter grown in cortical organoid differentiation medium (CODM). The colour scheme represents the value of -log_10_(P), where P is the probability of enrichment adjusted by a false discovery rate of 5%. The size of each circle represents the ratio of enrichment. Absent circles indicate an enrichment ratio < 1.1. **b**. The top five biological process synaptic terms not already shown in **a. c**. The top five biological process development terms not already shown in **a. d**. Comparison of the KEGG pathway terms. **e**. Examples of proteins detected exclusively in either iPSCs or cortical organoids involved in maintenance of stems cells or synaptic functions.

### 2D/3D iPSC-derived Cortical Brain Organoids showed mature neuronal morphology and function

Next, we sought to evaluate maturation and functionality of cortical organoids. Maturation of neurons is usually attributed to the presence of synapses. We first used IHC to examine synapse formation in organoids. MAP2 positive glutamatergic excitatory neurons expressing the vesicular glutamate transporter 1 (VGLUT1) protein were evident in brain organoids **(Figure 5a, b** (high magnification of boxed area in A)). Pre-synaptic protein synaptophysin was readily detected in puncta that localised in proximity to postsynaptic density protein PSD95 **(Figure 5c**). Similarly, inhibitory GABA neurons and a punctate pattern of presynaptic VGLUT1 were also evident in 2D cultures of dissociated of brain organoids (**Figure S4**). To comprehensively characterise the synaptic machinery in these organoids we analysed the proteome of 5 months-old organoids (n=3 organoids per batch, N=3 different batches of differentiation): cell type specific enrichment confirmed the presence of all expected cell types including neurons, astrocytes and oligodendrocytes. Finally, ultrastructure analysis by transmission electron microscopy (TEM) demonstrated the presence of typical synaptic structures showing synaptic vesicles and electron-dense synaptic contact sites (**Figure 5d, e**, asterisks and arrowheads, respectively).

**Figure 5.**
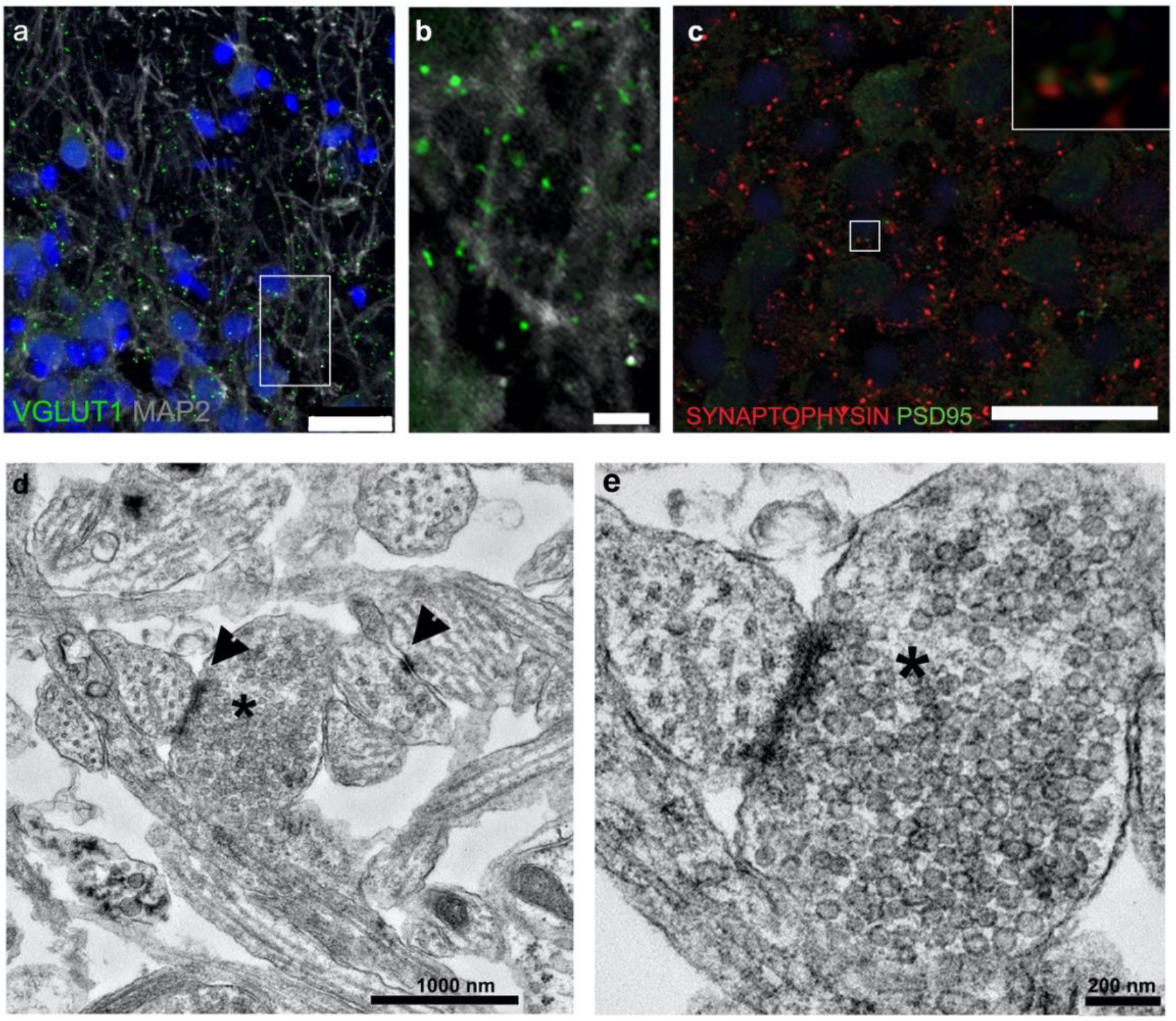
Cortical organoids synaptic maturation. **a, b**. Representative immunohistochemistry images showing VGLUT1 puncta in MAP2 neuronal dendrites (**b**, high magnification image of inset in **a**). **c**. SYNAPTOPHYSIN pre-synaptic and PSD95 post-synaptic synaptic markers. **d, e**. Ultrastructure electron microscopy images of cortical organoids showing synaptic clefts (head arrows) and synaptic vesicles (asterisks). Scale bars, 5 μm (b), 25 μm (a, c).

We next tested the neurophysiological activity of organoids by multielectrode array (MEA). Five months-old organoids were cultured in standard conditions until 2 months prior to recordings and then with either cortical organoid differentiation medium (CODM) or a physiological relevant neuronal medium (BrainPhys) up until the MEA recordings. Organoids were placed on the MEA one day before recordings (**Figure 6a**). Spontaneous firing activity was observed in organoids cultured in BrainPhys (**Figure 6b, c**), but not in CODM organoids. Similarly, spike raster plots showed firing patterns of organoids across all electrodes with marked network bursts in organoids cultured in BrainPhys only (**Figure 6d, e and S5a**). The mean firing rate (MFR) of neurons cultured in BrainPhys was significantly greater than CODM grown organoids (1.7 ± 1.3 Hz in BrainPhys and 0.03 ± 0.04 Hz in CODM (n = 5 organoids, N=3 independent batches of differentiation, mean ± SD 60 electrodes, paired two tailed t-test, p= 0.0466) (**Figure S5e**). Next, we performed pharmacological intervention using tetrodotoxin (TTX) as a synaptic blocker. Organoid network activity was abolished with TTX addition with activity returning to normal levels after washout (**Figure 6f**, n=5 organoids, N= 3 differentiation batches, mean ± SD, paired two tailed t-test, **** p<0.0001).

**Figure 6.**
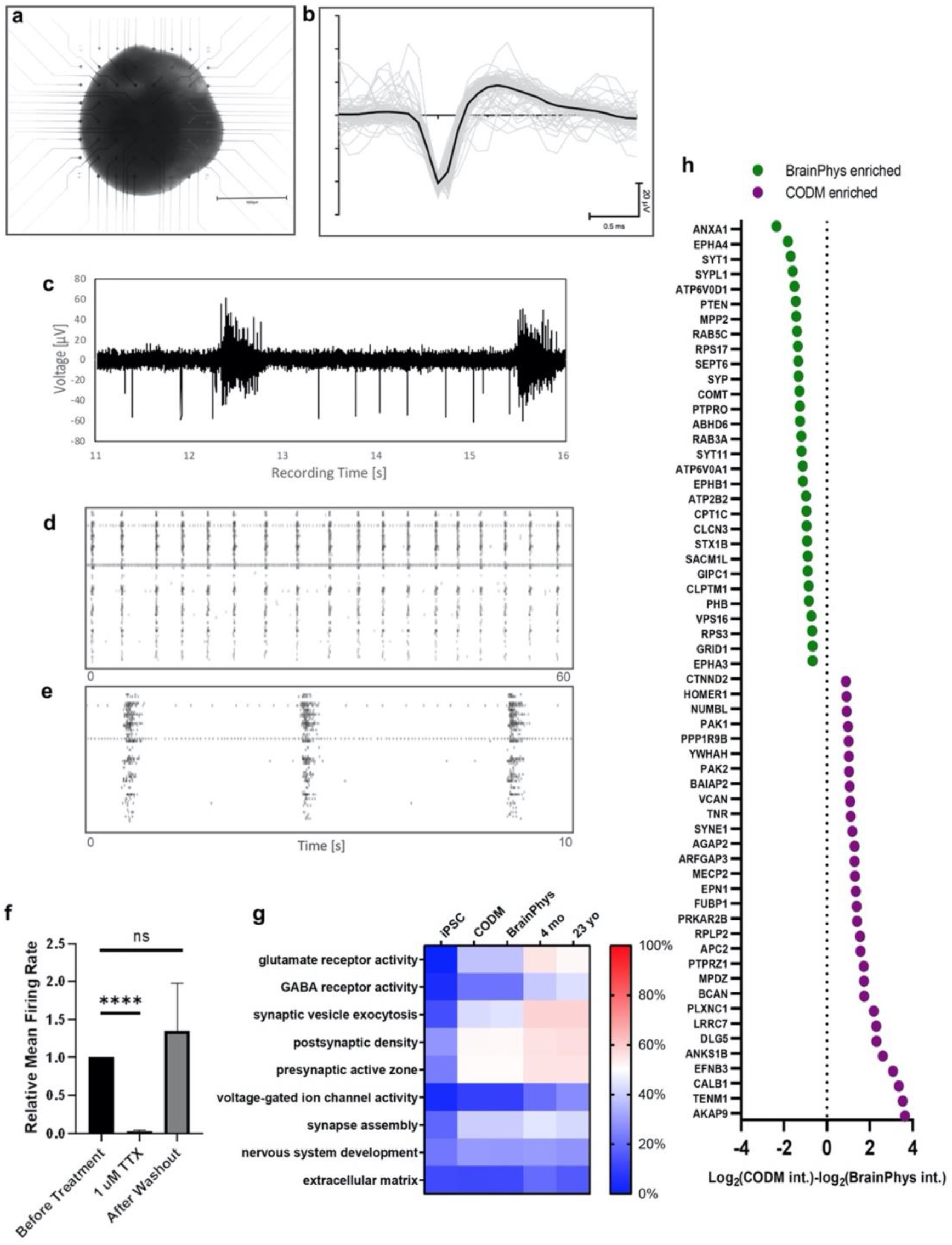
Cortical organoids develop synchronous networks. **a**. Image showing cortical organoid plated overnight on Multielectrode array (MEA). **b, c**. Representative spontaneous firing activity of organoid. **d, e**. Spike raster plots showing firing patterns of organoids across all electrodes with marked network bursts. **f**. TTX treatment abolished firing activity, which returned following (mean ± SD, paired two tailed *t* test, **** p<0.0001). **g**. Graph showing proteins significantly differentially enriched in organoids cultured in CODM or BrainPhys media (p < 0.05, adjusted by a false discovery rate of 5%).

BrainPhys medium has been described as a more physiological environment supporting the basic function of neurons^22^. To gain insights into possible differences between BrainPhys and CODM cultured organoids we compared the proteome of organoids cultured in the two media (**Figure 6g, h**, n= 3 organoids for each condition). Proteomic analysis detected 4,334 in common proteins for both conditions with an additional 27 and 242 unique Brain Phys and CODM proteins, respectively. The different culture conditions showed differences in protein enrichment (**Figure 6g**). Notably, enriched proteins that could potentially explain the improved function of BrainPhys cultured neurons included synaptic vesicle-associated proteins synaptophysin (encoded by *SYP*) and synaptotagmin 1 (encoded by *SYT1*), glutamate receptor delta 1 subunit (encoded by *GRID1*) involved in synaptogenesis and phosphatase and tensin homolog (encoded by *PTEN*) responsible for synapse maturation. Furthermore, organoids were transduced with an AAV9.SYN1 (*Synapsin 1*) promoter driving an mcherry reporter Synapsin 1 tethers pools of synaptic vesicles in nerve terminals, binds to the cytoskeleton, and has an activity-dependent function regulating the availability of synaptic vesicles for neurotransmitter release^24^. When cultured in BrainPhys, organoids showed increased neuronal *Synapsin 1* promoter activity when compared to standard CODM organoids (**Figure S5a, b**). We also confirmed the presence of synapses and synaptic vesicles in BrainPhys organoids using TEM (**Figure S5f**).

### 3D retinal-cortical organoids form optic nerve-like structures

Having demonstrated the formation of dorsal cortical organoids from 2D/3D confluent differentiation cultures, we next aimed to test whether the 3D suspension culture of complex organoids comprising both retinal and brain organoids promoted the formation of connections between the two organs.

In early 2D differentiation cultures, retinal and brain organoids spontaneously developed in proximity enabling the manual isolation of these two organoids for further neural maturation in suspension (**Figure 7a**). This differentiation potential was tested in 4 PSC lines (see Methods for details), including the H9.mcherry embryonic stem cell (ESC) line where the typical morphology of brain neural rosettes and thick neural epithelium of retinal organoids are easily discerned. At this early time point in 3D culture (5 weeks) the two organoids were attached but failed to show any clear axonal projections (**Figure 7b**). The connection of the retina to the brain to process the visual information is established by retinal ganglion cells (RGCs), the first-born cell type of the retina, which form neuronal outputs connecting to the brain through the optic nerve. In the complex retinal-brain organoids, at 10 weeks of development, the maturation of each individual organoid was evident. In retinal organoids NEUN and HuC/HuD colocalised with THY1 and MAP2 RGC markers delineating these cells’ axonal projections towards the centre of the organoid (**Figure S6a, b**). Similarly, these two markers were also expressed in neurons within the brain organoids, as demonstrated by the colocalization with TBR1 cortical neural marker (**Figure S6c, d**). CRX positive photoreceptor cells were present in retinal organoids with RGC THY1 positive axons clearly extending long axonal processes that connect to the TBR1 positive brain organoid (**Figure 6c, d**, n= 13 organoids, N= 9 differentiation batches). Finally, we used Lightsheet microscopy to image the morphological intricacies of whole 3D retinal-brain complex organoids (n=3 organoids, N=2 differentiation batches). Cleared organoids enabled the visualisation of MAP2 positive projections connecting both retinal and brain regions forming structures that resemble the optic nerve (**Figure 7e, h and SMovie 3**, n=3 organoids imaged).

**Figure 7.**
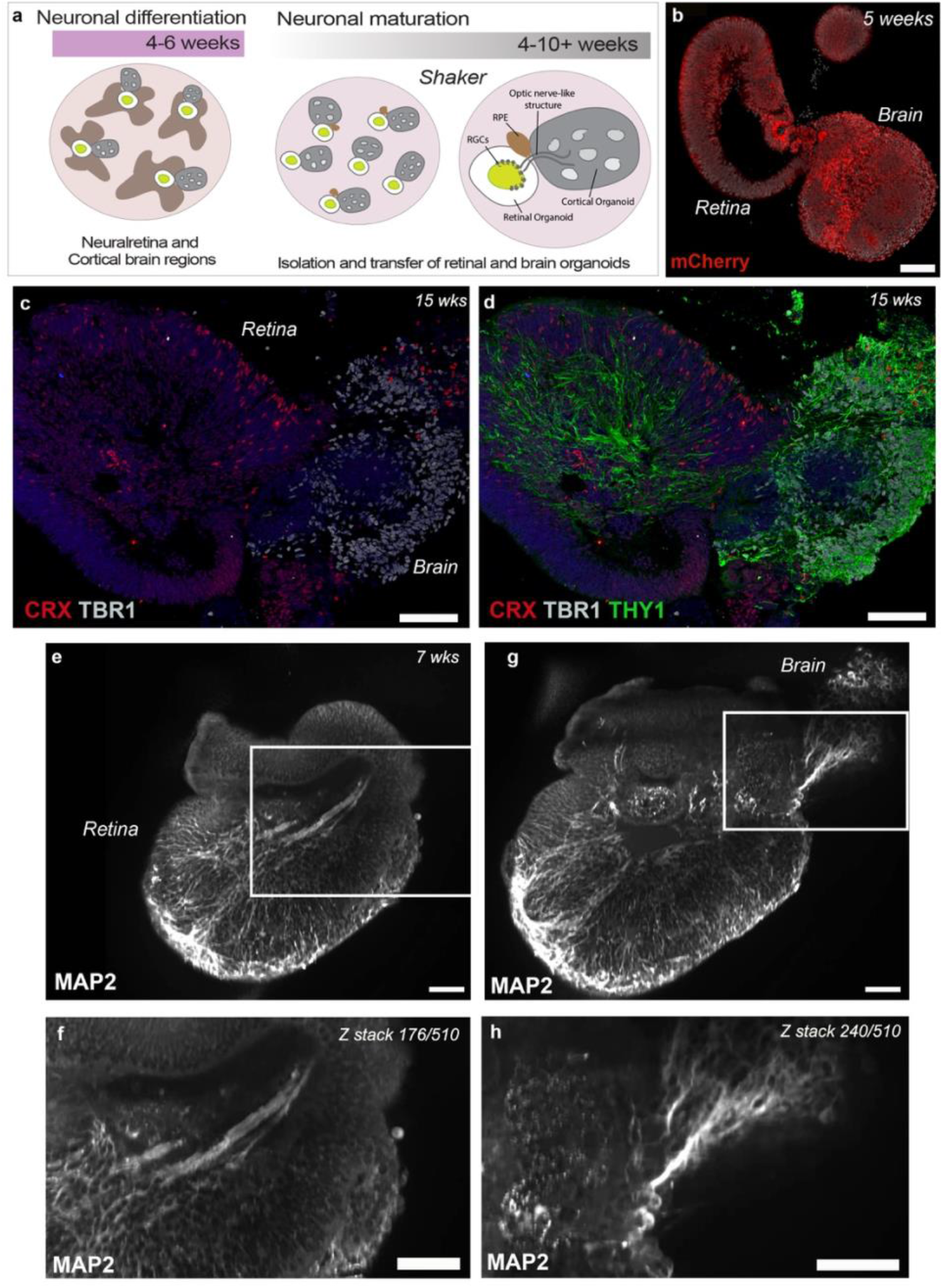
Complex retinal-cortical organoids form optic nerve-like structures. **a**. Schematic illustrating the timeline of neuronal differentiation and the main cellular components of the retinal-cortical system. **b**. Week 5 mCherry retinal-brain organoid showing retinal and brain regions growing together. **c**. Representative immunohistochemistry images of CRX positive photoreceptor cells in retinal organoids and TBR1 neurons in brain organoids. **d**. THY1 positive retinal ganglion cells send axonal projections to the centre of the retinal organoid that connect with brain organoid THY1 positive neurons in brain regions. **e-h**. Light sheet image of whole retinal-brain organoid showing consecutive z-stack images of MAP2 retinal ganglion cells axons. **e**. Forming optic nerve-like structure projecting towards the central part of retinal organoid (**e**). **f**. High magnification image optic nerve-like structure (z stack 176 out of 510 stacks) from inset in **g**. Deeper z-stack 240 showing the same axonal projection connecting to brain region. **h**. High magnification image of inset in **g** showing optic nerve-like structure. Scale bars, 100 μm (b-d), 200 μm (e-h).

## Discussion

The differentiation of PSCs into retinal and brain organoids have been used to model several neurodegenerations. A fundamental requirement for the development of therapies is the establishment of robust pre-clinical models in which they can be tested. Whilst numerous retinal and brain organoid differentiation protocols have been described, these differ considerably in methodology: some methods rely entirely on the addition of growth factors and matrices to guide cell fate decisions and organ formation while others form spontaneous organoids through the endogenous potential of PSCs to form the niches of miniaturized organs *in vitro*. Variability between protocols and, most importantly, in-batch organoid variability is a common problem of both approaches. Other challenges include the phenotypical and maturity differences between *in vitro* generated cells and their *in vivo* counterparts. Furthermore, the generation of the right cell type to enable the robust analysis of functional assays and testing of new therapies requires long-term cultures. Each of these protocols present their own advantages and disadvantages. However, to facilitate scale up and the high throughput use of these brain-like models, their variability and culture costing are usually the most common challenges and, thus, the initial choice of protocol is important.

Here, we adapted a previously-described simple and robust 2D/3D differentiation approach derived from confluent cultures of PSCs^19^ and generated dorsal cortical organoids within the same cultures of retinal vesicles and RPE cells. These cortical organoids are morphologically different and distinguishable from retinal vesicles which display thick neuroephithelia. Instead, brain organoids form the neural rosettes typical of cortical organoids, a feature that enabled their ready isolation. This differentiation method only requires a confluent culture of PSCs followed by change to a pro-neural induction media. Particularly important to reducing variability and costs of long-term cultures, this simple method does not require the addition of Matrigel or exogenous factors. Having tested this protocol in various PSC lines, we confirmed the generation of dorsal cortical organoids that express typical cortical plate markers as well as inhibitory and excitatory neurons and glial populations, with very few PSC lines failing to give rise to neuronal tissues.

The initial extracellular matrix and local cell-cell interactions provided by the 2D confluent environment of differentiating PSCs is sufficient to drive self-formation of discrete 3D structures, such as the eye and the brain. PSC cell lines vary in their efficiency to specify towards the three embryonic germ layers due to cell intrinsic cues specific to each cell line. Self-forming spontaneous methods benefit from the default bias present in most PSC lines to form ectoderm and forebrain^25^. Omics analysis of PSC lines and their initial stage of differentiation have shed light on the variability between cell lines leading to increased differences between organoids generated among multiple cell lines^10,26^. Our undirected protocol enables reproducibility of directed organoids but with the benefits of spontaneous differentiations.

Despite tremendous progress in the field, the lack of disease-relevant functional assays in organoids hinders their ability to test for new treatment efficacy. However, disease molecular signatures and biomarkers can be determined using integrative analysis of omics as well as computational or bioinformatic methodologies. In this study, cortical organoids generated by 2D/3D confluent method were extensively characterized using scRNA-seq and proteomics analysis, setting a baseline for future studies using disease organoids. In agreement with a previous correlation study comparing scRNA-seq datasets of brain organoids and foetal human brain^12^, our organoids showed a high correlation with brain organoids derived by other methodologies, particularly with dorsally patterned brain organoids generated in the directed differentiation approach demonstrated in Velasco et al., 2019^16^, known for their reproducibility.

The proteome of brain organoids has been seldom investigated. Previous proteomic analysis of early-staged (day 45) brain organoids generated using a whole-brain spontaneous 3D aggregation method highlighted the protein–protein interactions involved in early neuronal specification, including the migration of progenitor cells, radial glia, and oligodendrocyte progenitors^27^. Our proteomic analysis compared iPSCs and five months old cortical organoids. The analysis supported the increase in expression of proteins related to neuronal development and synaptic function in the cortical organoids and a relative decrease in cell cycle related proteins. The detection of synaptic protein components was further corroborated by IHC for synaptic markers and ultrastructure electron microscopy.

Neuronal network formation in brain organoids have previously been studied elegantly in late-stage organoids cultured on the MEA for a few months^10,15^ and in bioengineered neuronal organoids^28^. Our own electrophysiology analysis verified the functionality of cortical organoids, demonstrating their synchronous neuronal network activity when cultured for the final two months of differentiation in an electrophysiologically-relevant medium, BrainPhys^22^. A separate proteomics analysis identified proteins enriched in the BrainPhys-cultured organoids that might explain how BrainPhys media promotes favourable protein expression and electrophysiological activity. Overall, these results demonstrate the generation of functional cortical organoids that can be reliably differentiated for disease modelling and precision medicine studies.

Neurodevelopmental disorders leading to higher order cognitive abilities mostly affect the cortex and thus modelling of these conditions using cortical organoids is crucial^29^. In the eye, numerous retinal diseases have been investigated using retinal organoids^30,31-38^. However, modelling of more complex disease traits requires the differentiation of specific brain regions and their functional integration as well as their connection with other organs. One example of complex organotypic formation was demonstrated by the fusion of cortical, spinal cord organoids and skeletal muscle spheroids which generated a functional cortico-motor assembloid system leading to robust muscle contraction^39^.

In this study, we demonstrated the spontaneous formation of retinal-and-brain complex organoids in the 2D/3D confluent cultures. When in 3D suspension, these retinal-brain organoids maintained their proximity and were connected through an optic nerve-like structure mimicking the neuronal projections that connect the eye and brain. Our confluent method of differentiation enabled the precise isolation of these two organoids, overcoming variability within 3D directed and whole brain protocols that sporadically generate eye structures^8,10^ and the forced fusion of different organoids to form assembloids. The latter have been successfully used to model *in vivo* neuronal interactions between different brain regions^1,39-42^. The formation of optic nerve structures *in vitro* in complex organoid structures will better elucidate the dynamics of retinogenesis and neurogenesis that can then be used to model a number of optic neuropathies, such as glaucoma. Importantly, these retinal-brain organoids also promise to improve organoid development, such as the long-term survival and maturation of retinal ganglion cells in the retinal organoids as these cells would no longer lack their contact with the output in the cortex. Neurodegeneration in glaucoma is complex and extends beyond the eye into the lateral geniculate nucleus and visual cortex^18^ of the brain. Similarly, late onset neurodegenerative diseases, such Alzheimer’s show early phenotypes in the eye^17^, which enables pre-symptomatic diagnosis^43^. Accurate and more complex models of such diseases will elucidate novel aspects of disease pathogenesis and aid the challenges of developing effective treatments.

Our data demonstrate the simple and undirected derivation of both cortical and retinal organoids from a confluent culture of PSCs. The simplicity of this differentiation method coupled with the ease of precisely locating dorsal cortical organoids, due to their proximity to retinal vesicles, addresses the current problems of in-batch organoid-to-organoid variability. The comprehensive characterisation shown here, including novel proteome data of brain organoids, expands the available repertoire of relevant models of human disease. Importantly, this method also enabled the formation of complex organoid-forming structures that resemble the important connection, between the retina and the brain, the optic nerve. Future investigations of these neuronal populations and projections will allow the study of common optic nerve neuropathies.

## Materials and Methods

### Derivation of UCLOOi017-A-1 induced pluripotent stem cell (iPSC) line

Peripheral blood mononuclear cells (PBMCs) were isolated from the whole blood of a healthy donor using density gradient centrifugation. Briefly, 25ml of whole blood diluted 1:1 with phosphate buffered saline (PBS) was layered on top of 15ml of Ficoll-Paque Premium (GE) and centrifuged with brake and accelerator off at 500g for 30 minutes and the cloudy interphase containing PBMCs was collected. After washing once with PBS, the cells were counted and 2 million were cultured for 6 days in hematopoietic expansion media -Stemspan H3000(Stemcell Technologies), with the addition of EPO (R&D), IL-3 (Life Technologies), Dexamethasone (Sigma), Ascorbic Acid (Sigma), SCF (Miltenyi) and IGF-1 (Miltenyi). Following expansion, 200000 cells were nucleofected using Amaxa 4D nucleofector with Addgene plasmids pCXLE-hUL (Addgene plasmid #27080; http://n2t.net/addgene:27080; RRID:Addgene_27080), pCXLE-hSK (Addgene plasmid #27078; http://n2t.net/addgene:27078; RRID:Addgene_27078) and pCXLE-hOCT3/4-shp53-F (Addgene plasmid #27077; http://n2t.net/addgene:27077;RRID:Addgene_27077) [pCXLE-hUL, pCXLE-hSK and pCXLE-hOCT3/4-shp53-F were gifts from Shinya Yamanaka]. The nucleofected cells were plated on a well of a 6 well plate coated with Geltrex matrix (Life Technologies) and transitioned to Essential 8 media (Life Technologies)^1^. iPS cells were visible within a week and small colonies were visible within 2 weeks. Four to five weeks later, iPSC colonies were large enough to be excised, broken up and transferred using a 10µl pipette to individual wells of a 12 well plate (Corning). Once colonies had enlarged in the 12 well plate, each well (containing an iPSC clone) was dissociated and further cultured separately on 6 well plates (Corning) for 10 passages prior to characterisation.

### Human PSC maintenance

The human pluripotent stem cells including iPSC (UCLOOi017-A-1, ECCACs HPSI0214i-kucg_2 and HPSI0314i-hoik_1 and TiPSC-5^2^) and ESC (WiCell H9 WA09, H7 WA07) lines were maintained on feeder free conditions on Essential 8 media (E8, Life technologies) and Geltrex (Life technologies) coated 6 well plates. Briefly, when 70% confluent, PSCs were dissociated using Versene solution (Life technologies) at 37°C for 5 to 10 minutes. The dissociated clumps were washed once in PBS with centrifugation at 900 rpm for 5 minutes. The resulting cell pellet was broken up and resuspended in 1ml E8 media by 1000ul pipette and distributed on a Geltrex coated 6 well plate with E8 and 10µM ROCK inhibitor (Y-27632 dihydrochloride, Tocris) for 24 hours. Daily feeding with E8 was continued for further maintenance culture.

### Differentiation of PSC into cortical organoids

Human iPSCs were maintained until 90-95% confluent as described above. Media was replaced with Essential 6 media (E6, Life Technologies) for 2 consecutive days. At day 3 of differentiation, E6 media was replaced with a pro-neural induction media (PIM, composed of Advanced DMEM/F12, N2 supplement, L-Glutamine, non-essential amino acids and Antibiotic-antimycotic). At around week 3-4 of culture, three-dimensional (3D) organoids containing rosettes were observed throughout the plate and in close proximity to neuroretinal vesicles. The 3D cortical organoids were manually excised with 19G needles and kept together in 60 mm well plates in retinal differentiation media (RDM, composed of DMEM, F12 Nutrient mix, B27-vitamin A and antibiotic-antimycotic) and placed on an orbital shaker at 85 RPM. At 6 weeks of differentiation, retinal differentiation medium was supplemented with 10 % FBS, 100 μM Taurine (Sigma, T4871) and 2 mM Glutamax. At 10 weeks, cortical organoids were cultured in a cerebral organoid differentiation medium (CODM) as described in Lancaster & Knoblich, Nature Protocols, 2015 (Composed of Neurobasal medium, DMEM/F12, N2 supplement, insulin, Glutamax, MEM-NEAA, B-mercaptoethanol, B27 supplement) or BrainPhys™ hPSC Neuron Kit (Stemcell Technologies, composed of BrainPhys Neuronal medium, NeuroCult SM1 neuronal supplement, N2 supplement, human recombinant brain-derived neurotrophic factor, human recombinant glial cell line-derived neurotrophic factor, ascorbic acid and dibutyryl-cAMP). Differentiation cultures were fed every Monday, Wednesday and Friday. All representative images in the paper were from ***UCLOOi017-A-1*** iPSC line unless otherwise stated.

### Immunohistochemistry

hPSC-derived Cortical organoids were used for assessments of the time course of differentiation, as described in the main manuscript. Cortical Brain Organoids were washed with PBS, fixed for 40 minutes to 60 minutes depending on the size of the organoid in 4% paraformaldehyde and washed again with PBS prior to overnight suspension in 20% sucrose. The organoids, once sunken in sucrose, were embedded in OCT and snap frozen in nitrogen.

Cortical brain organoids were cryo-sectioned at 14 um thickness, collected on SuperFrost slides (Thermo) and preserved at minus 20°C. Cryosections were washed with PBS and blocked in 5% serum (goat or donkey) in blocking solution (1% Bovine serum albumin in PBS with 0.1% Triton-X) for 2 hours. Primary antibody (**Supplemental Table 1**) diluted in blocking solution was incubated overnight at 4°C. Sections were washed with PBS three times and incubated with secondary antibody (Alexa fluor 488, 546, 633 secondary antibodies, Invitrogen-Molecular Probes) diluted in blocking solution (1:500) at room temperature. Sections were then washed with PBS and counter stained with DAPI (Sigma-Aldrich). For immunohistochemistry of whole cortical organoids in 3D view using Lightsheet, a clearing protocol was performed. Briefly, cortical brain organoids were fixed in 4% paraformaldehyde, suspended in 20% sucrose and stored in 4°C as described above. Samples were incubated in clearing reagent 1 (Urea, N,N,N’,N’-tetrakis, Triton X, NaCl, dH_2_O) overnight followed by block in blocking solution (1% BSA in PBS, 5% serum and 0.3% Triton X) for 2 hours or overnight. Primary antibodies and DAPI were added in the blocking solution without serum overnight to increase penetration of antibody. Samples were washed with PBS and secondary antibodies diluted in blocking solution at 1:500 were added to the organoids overnight. The following day, samples were washed with PBS and clearing reagent 2 (Sucrose, Urea, Triethanolamine, Triton X, dH_2_O) was added overnight. Samples were embedded in low melting point agarose and left hanging in clearing reagent 2 for approximately 1 or 2 days until the samples were clear.

### Image acquisition

Images were acquired by confocal microscopy (LSM 880 Airyscan or SP5 Leica). A series of XY optical sections, approximately 1.0µm apart, throughout the depth of the section were taken and built into a stack to give a projection image. For Zeiss and Leica, ZEN blue and LAS AF image software were used, respectively. For Lightsheet fluorescence microscopy, images were acquired using Zeiss Lightsheet Z.1 using similar parameters to normal confocal using Zen Black software. Further image processing was performed in Imaris (version 9, Bitplane AG, Switzerland) software. Briefly, after images were imported, surfaces for each of the channels were created using the Surface Creation Wizard. Smoothing and threshold limits were applied to eliminate background light form the image. Optical sections of the organoid were created using the Oblique Slicer tool.

### Dissociation of Brain organoids

To produce 2D neuron cultures, cortical organoids were washed with PBS once and incubated in trypsin or Accutase for 5 minutes at 37°C. Organoids were then dissociated into single cells by gently pipetting up and down. Cells were then centrifuged at 300 *g* for 5 minutes and cell pellets were resuspended in 250 µl of media and seeded onto Geltrex coated 8 well chamber slides. Media was changed every other day until dissociated cells reached 11 weeks in culture.

For scRNA-seq, cortical organoids were dissociated with Neurosphere dissociation kit (Miltenyi Biotec). Cortical organoids were transferred to Eppendorf tubes then washed with 1 ml of PBS. Enzyme mix was prepared as per manufacture instructions and 500 ul added to each tube and incubated at 37°C for 10 minutes and flicked every 2 minutes. Organoids were then pipetted up and down with p1000 pipette to mechanically dissociate the big clumps of cells. The tubes were returned to 37°C for another 5 minutes and further dissociation. Following this incubation, a p200 pipette was used to break up the clumps further and 500 µl of HBSS was added to each tube and the contents were filtered using MACS 30 um filters into fresh Eppendorf tubes. Cells were pelleted by centrifugation at 400 *g* for 10 minutes at room temperature and resuspended in 200 µl of CODM media with the tubes kept on ice when transferred for cell counting.

### Quantification

Chamber slides were fixed by incubating in 4% PFA for 10 minutes and stained with TUJ1 and GFAP antibodies using the immunohistochemistry protocol described above. Next, cells were imaged using the Zeiss Axio Imager Alpha confocal microscope. Cell clumps cells were located using epifluorescence illumination and TUJ1 and GFAP positive cells were imaged from 10 different regions in the chamber slide and quantified using FiJi software. To accurately measure the percentage of TUJ1 neurons and GFAP glial cells, the number of DAPI, TUJ1 and GFAP cells were counted using cell counter plug-in. Then the percentages of TUJ1 and GFAP cells were calculated by in relation to the total number of DAPI cells.

### Single Cell RNA-Sequencing

Organoids were dissociated as described above and a subset of cell suspension was stained with 0.4% Trypan Blue (Gibco) and assessed for viability and concentration using Countess II Automated Cell Counter (Invitrogen). Single cell suspensions were passed through 40µm cell strainer (Corning) and concentration was adjusted to 1000 cells/µl. The suspension was loaded in single-cell-B Chip (10X Genomics) for target output of 10,000 cells per sample. Single-cell droplet capture was performed on the Chromium Controller (10X Genomics). cDNA library preparation was performed in accordance with the Single-Cell 3’ v3 protocol. Libraries were evaluated for fragment size and concentration using Agilent HSD5000 ScreenTape System. Samples were sequenced on an Illumina NovaSeq6000 instrument according to manufacturer’s instructions (Illumina). Sequencing was carried out using 2×150 paired-end (PE) configuration with a sequencing depth of 40,000 reads per cell. The sequences were processed by GENEWIZ. The data supporting the findings in this publication have been deposited in NCBI’s Gene Expression Omnibus and are accessible through GEO Series accession number GSE174232 (https://www.ncbi.nlm.nih.gov/geo/query/acc.cgi?acc=GSE174232).

### Single Cell RNA Sequencing analysis

Raw files were processed with Cell Ranger 2.0.1 software (10X Genomics). The Cell Ranger count module was used to align sequence reads from scRNA-seq experiment to the GRCh38 human reference genome and to generate the cell-by-gene count matrix. Genes and cells were filtered retaining those that are expressed in at least three cells and those that have at least 500 genes detected. Cells were further filtered by removing (i) those that have over 10 percent mitochondrial genes in all detected genes and (ii) predicted doublets using DoubletFinder (Version 2.0.3)^3^ with an expected rate of 2.5% (according to 10X Genomics guidelines). To compare our cortical organoid scRNA-seq data with other published datasets, a combined cell-by-gene count matrix of the processed scRNA-seq datasets with cell annotation was obtained from OrganoidAtlas (https://cells-test.gi.ucsc.edu/?ds=organoidatlas)^4^. We subset six scRNA-seq datasets from six different brain organoid studies each using a different culture/differentiation protocol. These include Giandomenico et al., 2019 (denoted as “Gian 3m U”), Birey et al., 2017^5^ (Birey 3m D), Velasco et al., 2019^6^ (Vela 3m D), Quadrato et al., 2017^7^ (Quad 3m U), Trujillo et al., 2019^8^ (Truj 6m D), and Velasco et al., 2019^6^ (Vela 6m D). A fetal brain dataset is also included for comparison (Zhong et al., 2018)^9^ (Fetal). The cell type compositions of each public dataset were visualized as staked bar plots. Cell annotation of the cortical organoid scRNA-seq generated in this study was carried out by using scClassify (Version 1.0.0)^10^. Briefly, we trained prediction models using each published dataset as reference separately and joint trained. Then, we predicted the cell types in our cortical organoid scRNA-seq based on the individual and the joint prediction models. A t-distributed stochastic neighbour embedding (t-SNE) was applied to visualize the cells with labels generated from the joint prediction model. To compare agreement in cell-type composition between our cortical organoid and the public brain organoid and fetal brain datasets, intraclass correlation coefficient (ICC) was calculated by generating a table of cell-type composition across published organoids and our organoid, using the ICC function in the irr R package (Version 0.84.1). To evaluate the reproducibility of cell type prediction, logarithm transformed data were used for assessing the expression of a panel of marker genes listed in^4^ in each cell type based on the scClassify prediction. The marker genes include neuronal growth cone markers (STMN2, GAP43, DCX), early neurogenesis markers (VIM, HES1, SOX2), excitatory neuron markers (TBR1, SLC17A7), inhibitory neuron markers (GAD1, SLC32A1), radial glial marker (EOMES), cell cycle markers (TOP2A, MKI67), oligodendrocyte precursor cells markers (OLIG1, OLIG2), astrocyte markers (S100B, SLC1A3, GFAP), cilium markers (MNS1, NPHP1), BMP signal markers (BMP4, MSX1), mesoderm marker (MYH3), proteoglycan markers (BGN, DCN) and unfolded protein response cell marker (DDIT3).

### Electron microscopy (EM)

Brain organoids were fixed overnight in Karnovsky’s fixative (2.5% glutaraldehyde and 2.4% formaldehyde in MOPS buffer), then washed in MOPs buffer. They were post fixed with 2% osmium tetroxide, incubated in 2% aqueous uranyl acetate, dehydrated in an ethanol series, and embedded in TAAB Low Viscosity Resin (TAAB Laboratories). Sections were cut at 90nm using a UC6 ultramicrotome (Leica Microsystems), then stained with 2% uranyl acetate in 50% ethanol (10 min) and Reynold’s lead citrate (4 min). Grids were examined with a JEOL JEM-1400 transmission electron microscope operating at 80 kV, with images recorded using a Matataki Flash sCMOS camera.

### Recording of organoid activity on Multiple Electrode Array (MEA)

After media replacement as described previously, a single organoid for each recording was placed onto a 60-electrode MEA200/30iR-Ti multiple electrode array (MEA) plate (MultiChannel Systems), ensuring coverage of the electrodes. Medium was removed and 20 μl of Matrigel extracellular matrix (Corning) was dropped onto the organoid. The MEA plate was subsequently incubated at 37 °C for 15 mins to allow the Matrigel to set the organoid in place. 500 μl of medium was added dropwise and the plate was placed back in the incubator at [37 °C, 5% CO2, 70% humidity] for 24 hours before recording electrical activity.

Organoid electrical activity was measured using an MEA2100-lite system with TC01 temperature control (MultiChannel Systems), heated to 37 °C. Recordings were made for 5-10 minutes at a frequency of 10 kHz using MultiChannel Experimenter software (MultiChannel Systems). Data was processed using the MultiChannel Analyzer software (MultiChannel Systems). Raw electrode recordings were filtered sequentially with a 200 Hz second order Butterworth high pass filter then a 3 kHz second order Butterworth low pass filter. Spikes were picked using a threshold of 5 standard deviations below mean voltage for each electrode.

Voltage-gated sodium channel activity was blocked using Tetrodotoxin (TTX). Briefly, baseline activity was recorded by MEA as described above. Immediately afterwards, Brainphys media was removed, and washed once with PBS, then replaced with Brainphys media containing 1µM TTX. Following a 10 minute incubation in culture conditions, a further MEA recording was taken. Subsequently, the media containing TTX was removed, and the organoid was washed 3 times with PBS, with fresh Brainphys media added after the final wash. Following 2 hours in the incubator, a final recording was taken.

Results were produced by counting the spikes using a -5SD threshold after filtering, deriving an average spike number for the whole MEA, deriving an average frequency for the whole MEA (by dividing the spike number by recording time), and normalising each organoid’s pre-, during and post-treatment frequencies to the frequency of the organoid during the pre-treatment recording. Normalised values were assessed for significance independently using a paired t-test, assuming similar distributions between the two datasets.

### Preparation of organoids and analysis by mass spectrometry

Six organoids were isolated and cultured as previously described until week 10 after initiation of differentiation. Three of these organoids were either cultured in CODM or BrainPhys medium for two weeks. Medium was removed from organoids which were then homogenised and lysed with a drill and pestle in 200 μl 2% SDS Lysis Buffer [2% (w/v) sodium dodecyl sulfate (SDS), 50 mM HEPES pH 7.4, 2 mM ethylene glycol-bis (β-aminoethyl ether)-N,N,N′,N′-tetraacetic acid (EGTA), 2 mM ethylenediaminetetraacetic acid (EDTA), 2 mM phenylmethylsulfonyl fluoride (PMSF), EDTA-free Protease Inhibitor (Roche) and PhosSTOP (Roche)] and freezing on dry ice. Frozen organoid lysate was thawed and reduction was performed with 10 mM tris(2-carboxyethyl)phosphine (TCEP) at 85 °C for 10 min with shaking. Samples were alkylated with 20 mM iodoacetamide for 30 min at 23 °C in the dark. Protein was precipitated from using the chloroform-methanol method^11^. Protein pellet was reconstituted in 20 μL of 7.8 M Urea, 50 mM HEPES pH 8.0. Protein was digested by the addition of 3 μg Lys-C (FUJIFILM Wako Pure Chemical Corporation) and incubation for 8 hours at 25 °C, 900 rpm. Sample was then diluted 8-fold with 50 mM HEPES pH 8.0 and digested by the addition of 5 μg TrypZean recombinant trypsin (Sigma-Aldrich) and incubation for 8 hours at 30 °C, with shaking. The trypsin digestion was repeated for each sample. Protein content of each digest was estimated by measuring the absorption of 280 nm light (Implen Nanophotometer, Labgear, Australia). 50 μg aliquots of each sample were desalted on an in-house made STAGE tip with C18 material. Samples were eluted in 50% acetonitrile, dried and then reconstituted in a solution of 90% acetonitrile, 0.1% trifluoroacetic acid (TFA) for hydrophilic interaction chromatography (HILIC) fractionation.

HILIC fractionation was performed on a Dionex Ultimate 3000 HPLC system with a 250 mm long and 1 mm inside diameter TSKgel Amide-80 column (Tosoh Biosciences). HILIC gradient was between a solution of 90% acetonitrile, 0.1% TFA (Buffer A) and a solution of 0.1% TFA (Buffer B). The sample was injected into a 250 μl sample loop to be loaded onto the column at a flow rate of 60 μl/min in Buffer A for 10 min. Gradient was from 100% Buffer A to 60% Buffer A (balanced with Buffer B) for 35 min at a flow rate of 50 μl/min. Fractions were collected into a 96-well plate using a Probot (LC Packings) at 1-minute intervals, monitored by absorbance of UV at 214 nm. UV signal was used to combine select fractions into similar amounts of peptide. Fractions were dried and reconstituted in 0.1% formic acid for LC-MS/MS analysis.

The LC-MS/MS was performed using a Dionex UltiMate 3000 RSLC nano system and Q Exactive Plus hybrid quadrupole-orbitrap mass spectrometer (Thermo Fisher Scientific). The content of HILIC fractions was loaded directly onto an in-house 300 × 0.075 mm column packed with ReproSil Pur C18 AQ 1.9 μm resin (Dr Maisch, Germany). The column was heated to 50 °C using a column oven (PRSO-V1, Sonation lab solutions, Germany) integrated with the nano flex ion source with an electrospray operating at 2.3 kV. The S lens radio frequency level was 60 and capillary temperature was 250 °C. The 5 µL sample was injected into a 20 µl loop and loaded onto the column in 99% reversed phase buffer A (solution of 0.1% formic acid) and 1% buffer B (solution of 0.1% formic acid, 90% acetonitrile) for 25 min at 300 µL/min. The sample was loaded in 99% buffer A for 25 min. The gradient was from then from 99% buffer A to 95% buffer A in 1 min, to 75% buffer A in 75 min, to 35% buffer A in 8 min, to 1% buffer A in 1 min, held at 1% buffer A for 2 min, to 99% buffer A in 1 min and held for 10 min. MS acquisition was performed for the entire 120 min.

All samples and fractions were analysed using data-dependent acquisition LC-MS/MS. For data-dependent acquisition, the MS scans were at a resolution of 70,000 with an automatic gain control target of 1,000,000 for a maximum ion time of 100 ms from m/z 375 to 1500. The MS/MS scans were at a resolution of 35,000 with an automatic gain control target of 200,000 and maximum ion time of 115 ms. The loop count was 12, the isolation window was 1.2 m/z, the first mass was fixed at m/z 140 and the normalized collision energy was 28. Singly charged ions and those with charge >8 were excluded from MS/MS and dynamic exclusion was for 35 s.

The raw LC-MS/MS data was processed with MaxQuant v1.6.7.02^12^ using the following settings: variable modifications were oxidation (M), acetyl (protein N-terminus), deamidation (NQ); carbamidomethyl (C) was a fixed modification; digestion was set to specific for trypsin with a maximum of 3 missed cleavages; Label Free Quantification (LFQ) was enabled; Homo sapiens reference proteome with canonical and isoform sequences downloaded March 9 2020 was used as well as the inbuilt contaminants file; minimum peptide length was 7 and maximum peptide mass was 5000 Da; second peptides search was enabled while dependent peptides searches were disabled; peptide spectrum matching and protein false discovery rates were set at 1%; minimum score for modified peptides was 40; all modified peptides and counterpart non-modified peptides were excluded from protein quantification; matching between runs was disabled. All other parameters were default.

Entries in proteinGroups results file were filtered: CON_ and REV_ entries were removed; entries with < 2 unique + razor peptides were removed; entries with no Intensity values were removed; entries for which no gene name could be derived were removed. LFQ Intensity values for remaining entries was used as input for an R script that used limma statistics to compare protein relative intensity statistically.

The code for this script is reported here: https://github.com/ChildrensMedicalResearchInstitute/MS_intensity_diffexpress.

The mass spectrometry proteomics data have been deposited to the ProteomeXchange Consortium via the PRIDE^13^ partner repository with the dataset identifier PXD025933 and 10.6019/PXD025933.

### Gene ontology enrichment analysis of iPSC and cortical organoid proteomes

The list of proteins detected by mass spectrometry were filtered to remove proteins that were not detected in all three biological replicates. The number of “razor plus unique” peptides was required to be two or greater using an average value across the three biological replicates. The filtered protein list was converted to a gene list. Duplicate genes arising from multiple protein accession and isoforms were filtered by removed the gene with the least number of “razor plus unique” peptides. Over-representation analysis was conducted using WebGestalt. Significant enrichments for biological process and KEGG pathway terms were determined by comparing to the human coding genome. The probability of significant enrichment was adjusted using a false discovery rate of 5%.

### AAV transduction of brain organoids

AAV viral vectors (1-3.6×10E11 vg/organoid) were added to a total volume of 375 μl using fresh CODM media used to culture the cortical and whole brain organoids. The organoids were then transferred to low binding 24 well plates (Costar, Corning) and media was completely replaced with CODM containing the AAV vectors. Cortical and whole brain organoids were incubated at 37 °C for half a day before adding another 625 μl of fresh media. After overnight culture at 37 °C, the organoids and CODM/vector mixture were transferred to a 60 mm dish. The dish was topped up to 4 ml with fresh CODM media and put on an orbital shaker at 85 RPM at 37 °C. After 48 hours, organoids were fed every Monday, Wednesday and Friday.

### Statistical analysis

All means are presented as mean ± SD (standard deviation) unless otherwise stated; N, number of independent experiments i.e differentiation batches, proteome samples or MEA measurements; n, number of images or retinal organoids examined, where appropriate. Statistical differences between two groups were tested by two tailed paired and unpaired *t* tests. The preformed test is specified in figure legends and main text. Statistical significance was assessed using Graphpad Prism software. Figures were generated in Adobe Photoshop.

## Acknowledgments

We thank Dr. Scott Page and Joshua Studdert from the Children’s Medical Research Institute Advanced Microscopy Centre and the ACRF Telomere Analysis Centre supported by the Australian Cancer Research Foundation, for providing access to microscopy equipment. We thank Emma Kettle for Electron Microscopy, performed at the Westmead Scientific Platforms, which are supported by the Westmead Research Hub, the Cancer Institute New South Wales, the National Health and Medical Research Council and the Ian Potter Foundation. We thank Joshua Studdert and Hillary Knowles from the Single Cell Analytics Facility which is supported by NSW Luminesce Alliance. We acknowledge the support of the Children’s Medical Research Institute Biomedical Proteomics Facility. Cell line iPS TiPSC-5 was kindly provided by Prof Yvan Arsenijevic and Prof David Gamm. This work is supported by Luminesce Alliance (PPM1 K5116/RD274) - Innovation for Children’s Health for its contribution and support. Luminesce Alliance - Innovation for Children’s Health, is a not for profit cooperative joint venture between the Sydney Children’s Hospitals Network, the Children’s Medical Research Institute, and the Children’s Cancer Institute. It has been established with the support of the NSW Government to coordinate and integrate paediatric research. Luminesce Alliance is also affiliated with the University of Sydney and the University of New South Wales Sydney.

## Author Contributions

M.F. and S.L contributed to conception, design, execution and analysis of all experiments; J.R.W. contributed to proteomics and multielectrode array experiments; D.X and H.J.K contributed to transcriptome data analysis; G.C.S. contribute to tissue culture experiments and immunohistochemistry; R.R.A. provided UCLOOi017-A-1 iPS cell line and contributed to manuscript revision; P.Y. contributed to design and transcriptome data analysis and manuscript writing; M.E.G. contributed to conception, design, execution of proteomics and multi electrode array experiments and manuscript writing; A.G.C contributed to the conception, design, execution and analysis of all experiments, manuscript writing and funding.

## Supplementary information

### Supplementary Figures

**Supplementary Figure 1.**
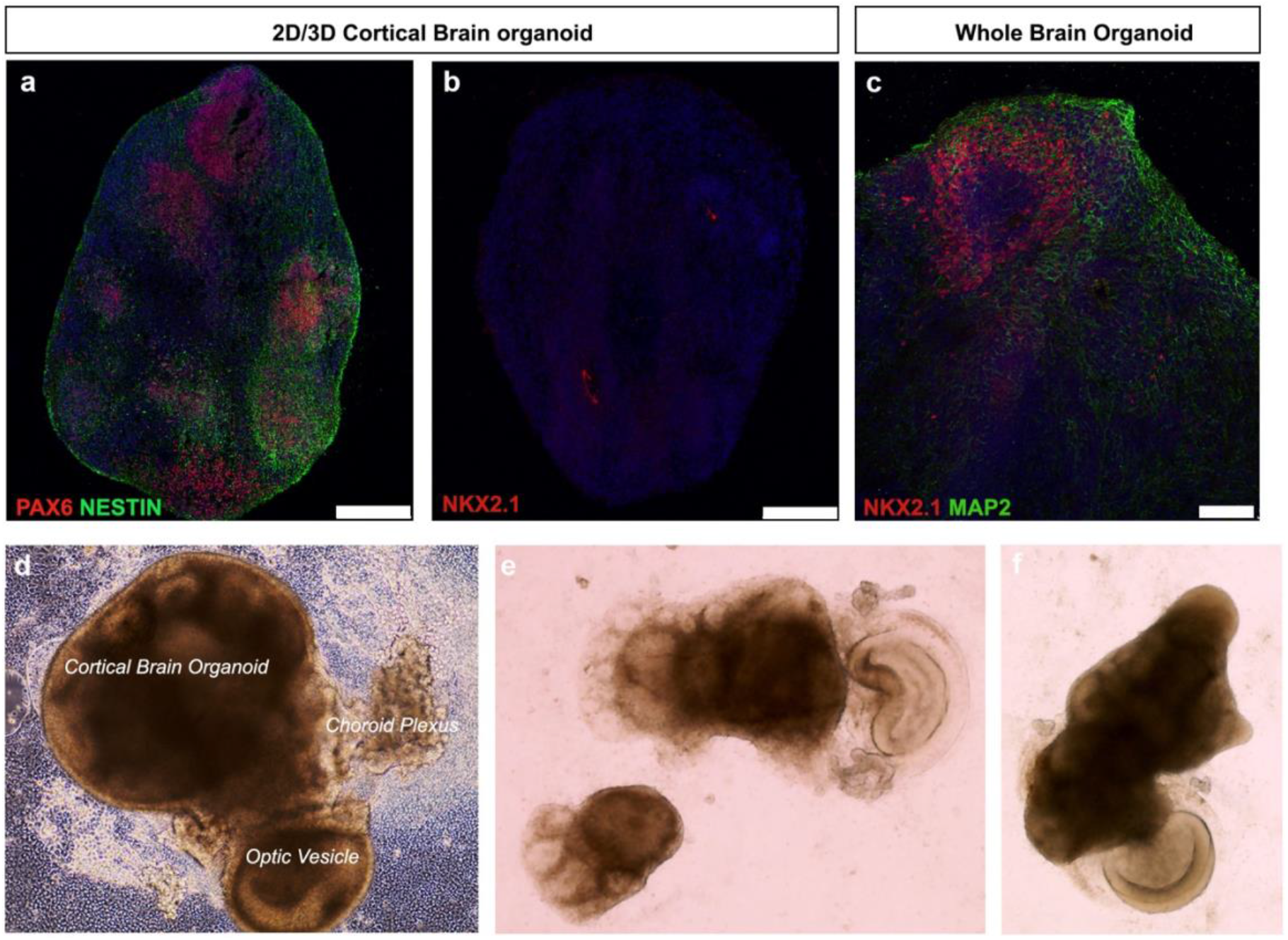
Confluent iPS differentiation protocol gives rise to cortical and retinal organoids. **a**. Immunohistochemistry image showing PAX6 and NESTIN positive cortical organoid. **b**. Image showing that cortical organoid negative for ventral cortex marker, NKX2.1. **c**. Image showing NKX2.1 and MAP2 positive whole brain organoid. **d-f**. Bright field imaged showing examples of cortical brain organoids growing in proximity to optic vesicles. Scale bars, 75 μm (c), 250 μm (a, b).

**Supplementary Figure 2.**
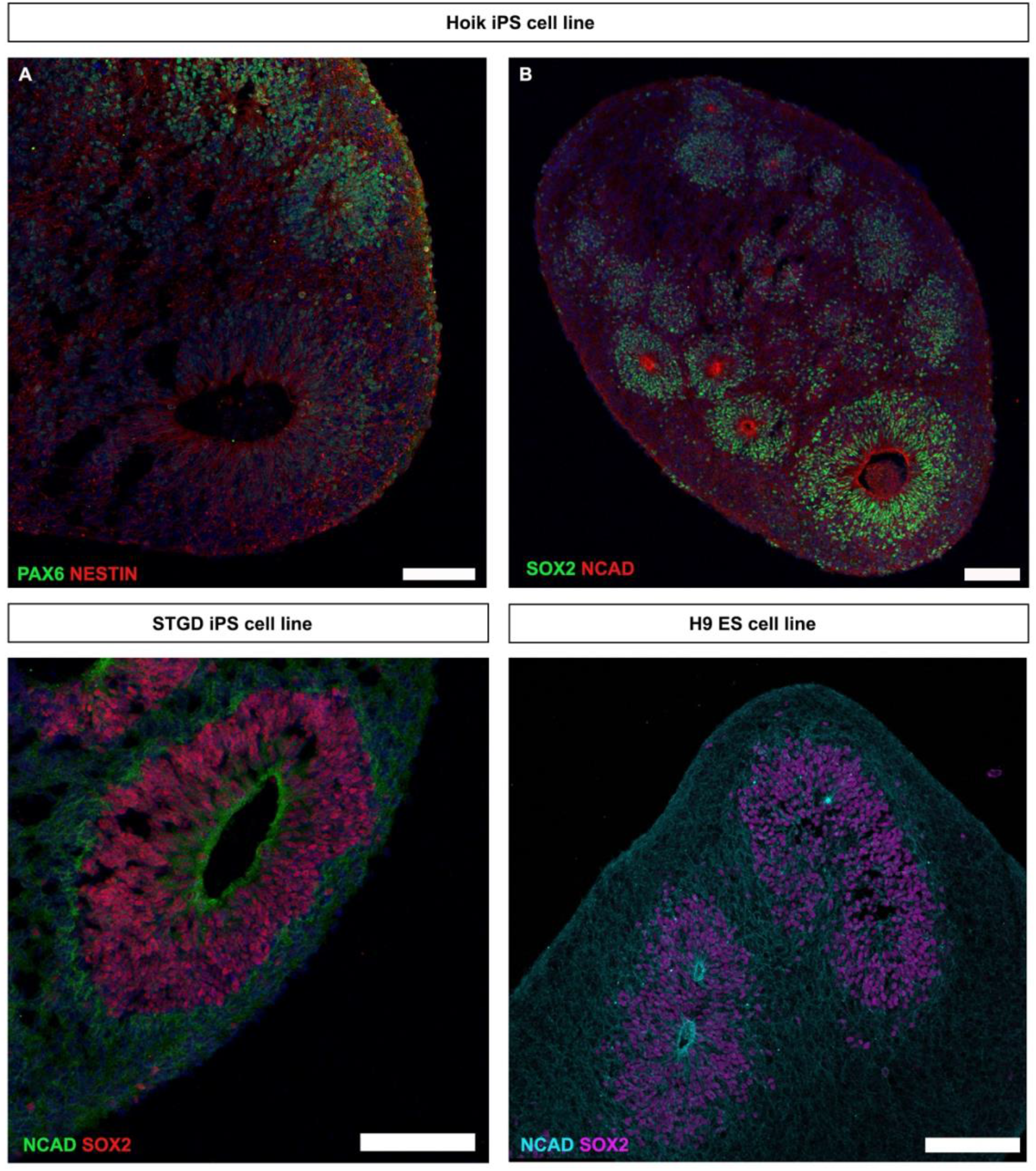
Early cortical organoids derived from various iPSC lines. **a-d**. Representative images of early neural markers of cortical organoids generated from iPSC lines and ES cell line H9. Scale bars, 100μm (a-d).

**Supplementary Figure 3.**
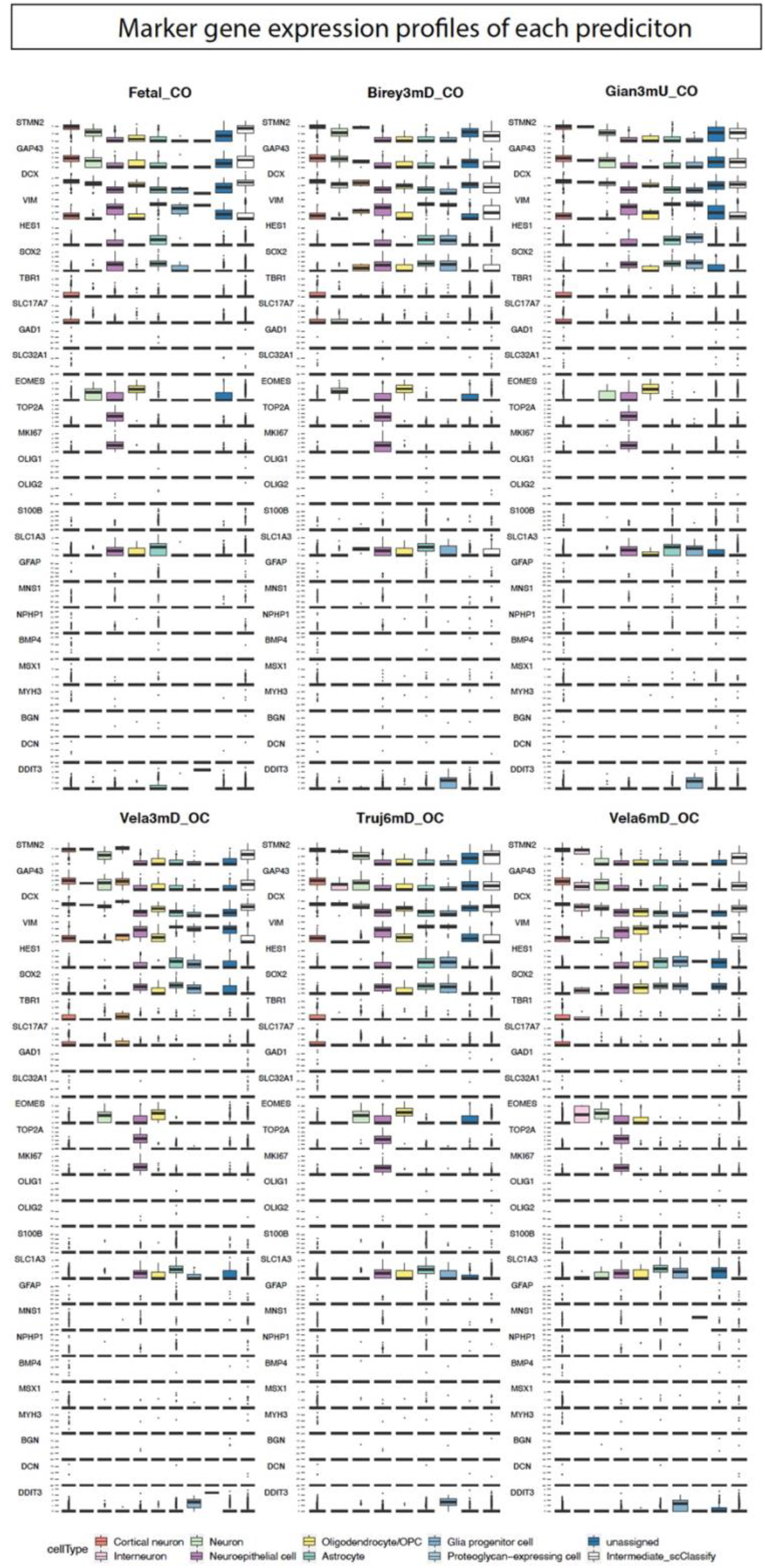
Marker gene expression profile for each scRNA-seq available datasets.

**Supplementary Figure 4.**
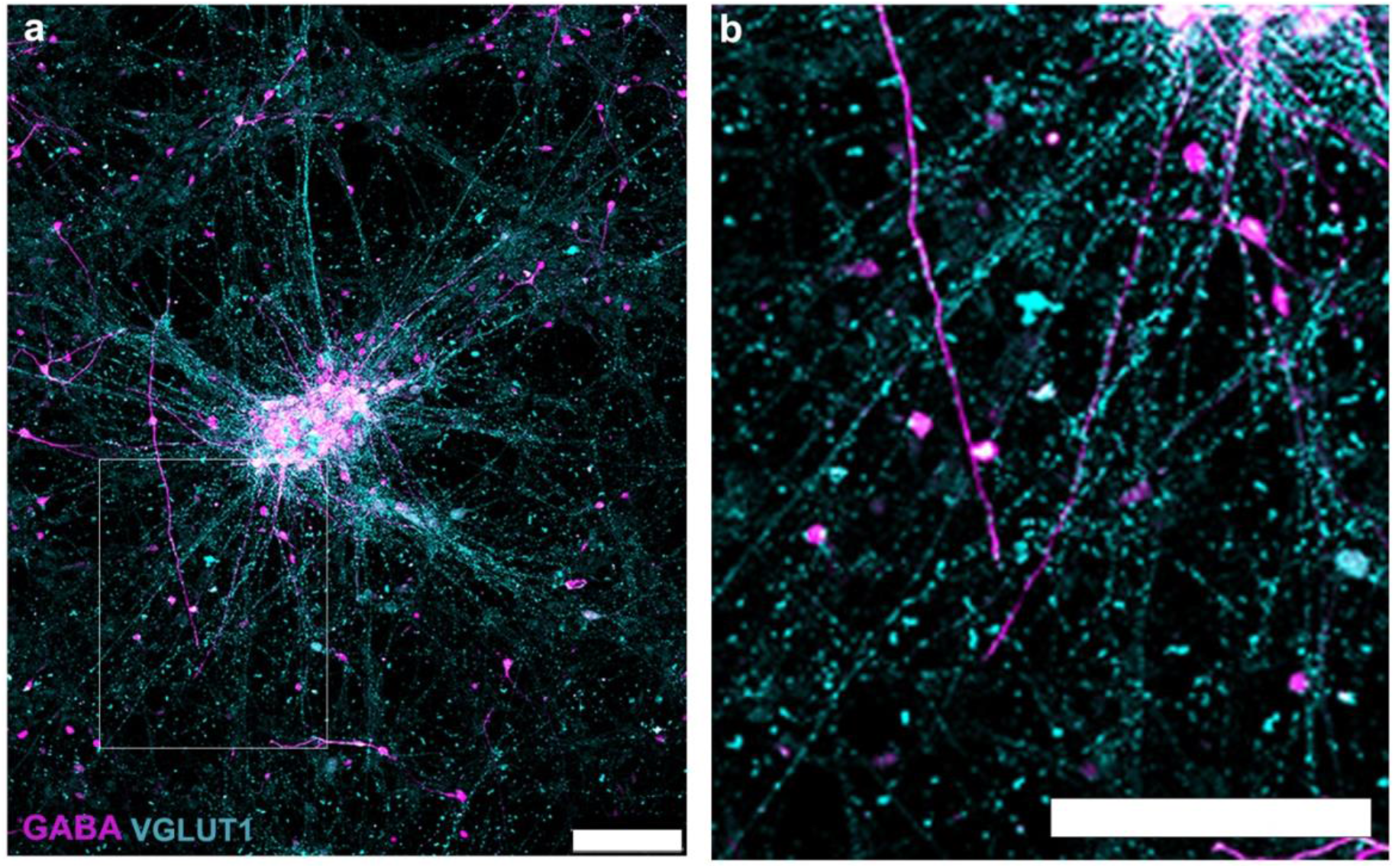
2D cultures of cortical neurons dissociated from brain organoids. **a, b**. Immunohistochemistry image showing GABA inhibitory neurons and VGLUT1 inhibitory neurons. b is the high magnification image of inset in a. Scale bars, 100 μm (a, b).

**Supplementary Figure 5.**
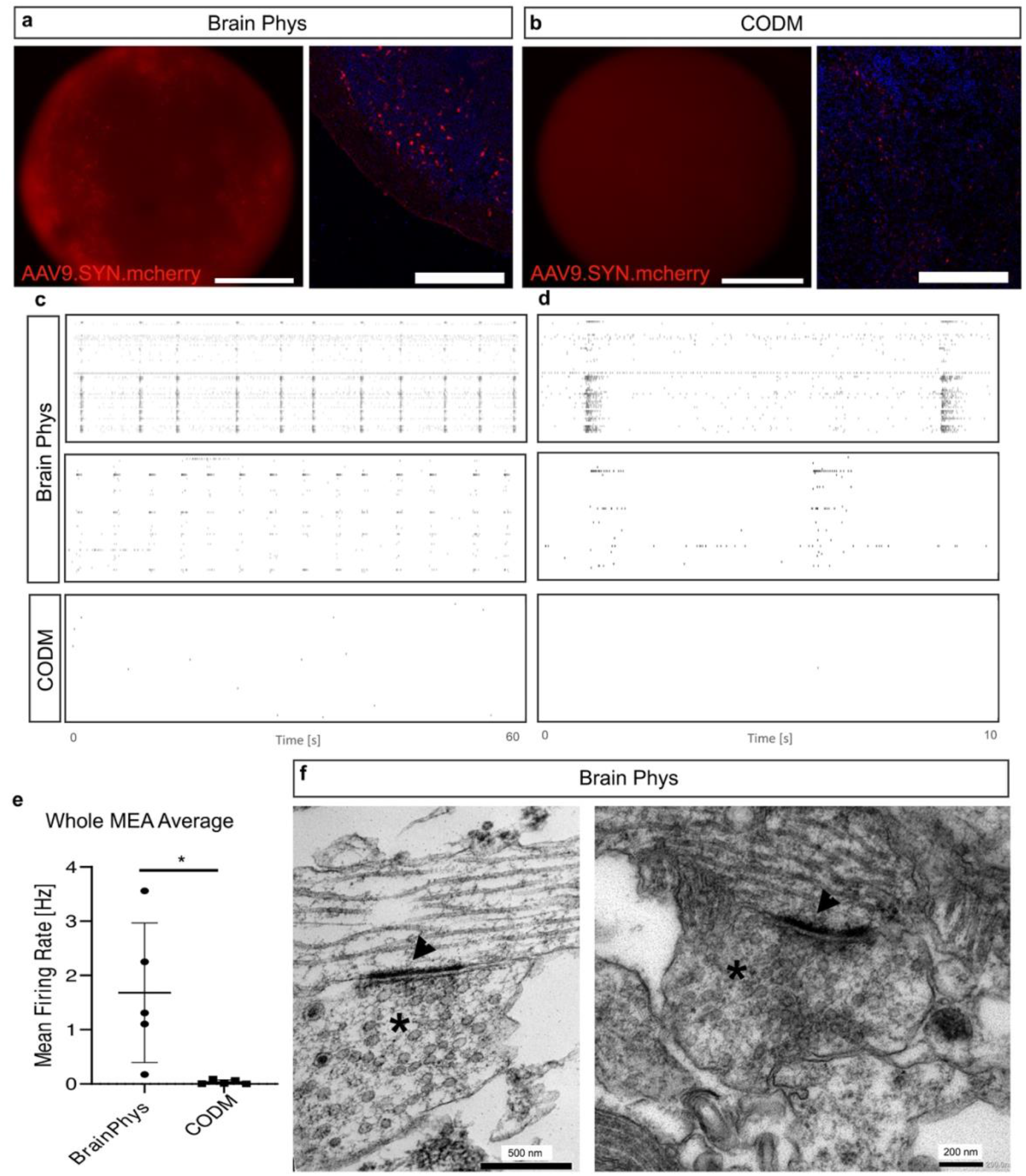
**a, b**. Representative bright field and immunohistochemistry images showing brain Phys and CODM cultured cortical organoids transduced with AAV9.SYN.mcherry viral vectors. **c, d**. Representative spike raster plots showing firing patterns of organoids across all electrodes with marked network bursts in Brain Phys, but not in CODM cultured organoids. **e**. whole MEA mean firing rate activity (mean ± SD, paired *t* test, * p<0.01). **f**. Ultrastructure electron microscopy images of Brain Phys cultured cortical organoid showing synaptic clefts (head arrows) and synaptic vesicles (asterisks). Scale bars, 250 μm (a, b)

**Supplementary Figure 6.**
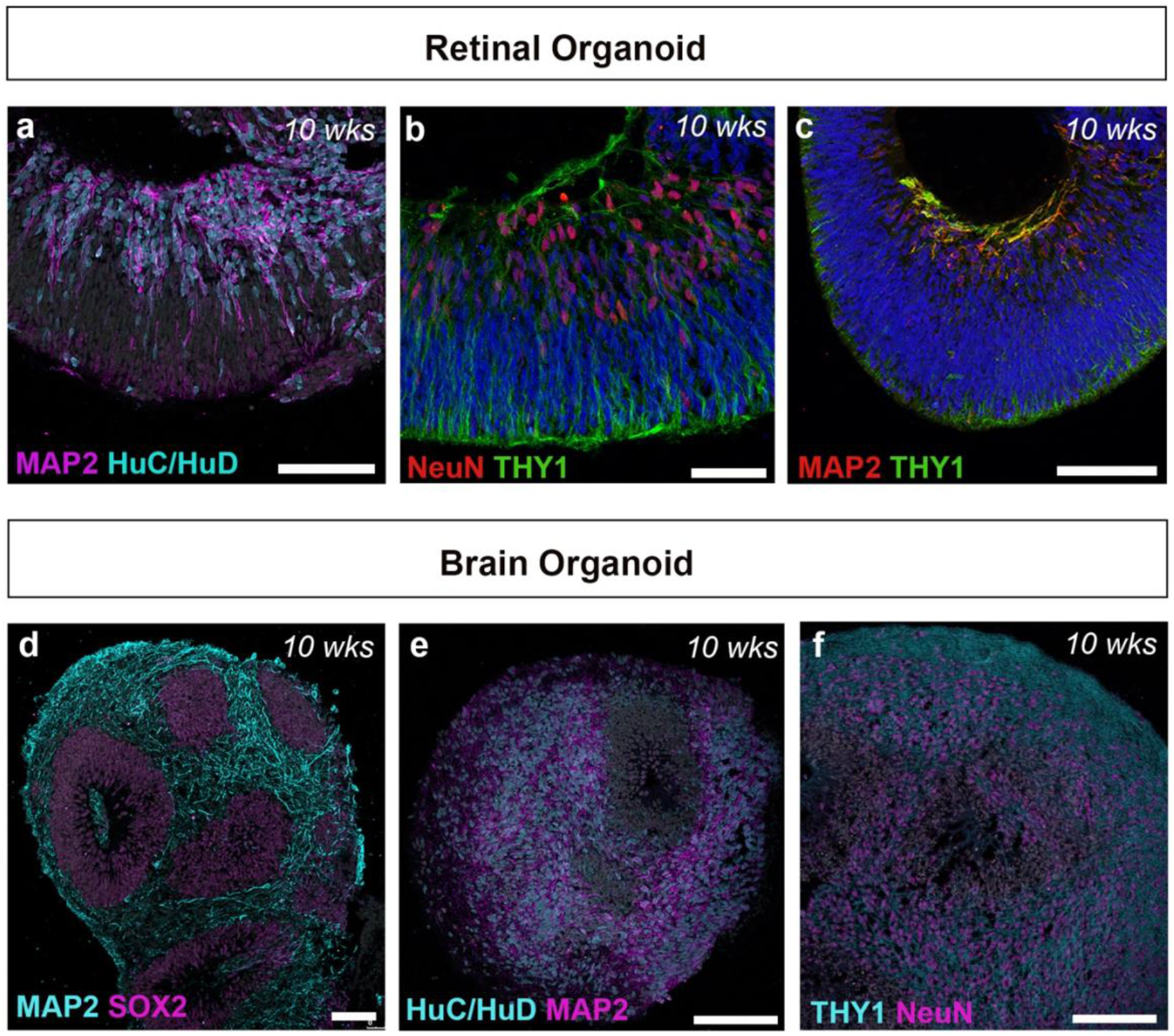
Neuronal markers on both retinal and brain organoids. **a-f**. Representative immunohistochemistry images from 10 weeks old retinal and cortical brain retinal organoids. a-c. Typical interneurons and retinal ganglion cell cells markers in the basal layer of the retinal organoid. **d-f**. The same markers are also expressed in cortical neurons of brain organoids. Scale bars, 75 μm (d), 150 μm (a, b, c, e, f).

### Supplementary Movies Legends

**SMovie 1. Differentiation of early cortical progenitors**

Video of a typical representative day 35 cortical organoid showing neuroepithelium regions positive for PAX6 (red) and NESTIN (green). DAPI stained in blue.

**SMovie 2. Differentiation of neurons and glial cells**

Lightsheet video of a cleared whole cortical organoid at day 75 in culture. Organoid showing numerous neurons positive for CALRETININ (red) and glial cells positive for SB100 (green). DAPI stained in blue.

**SMovie 3. Optic nerve-like formation between retinal and cortical organoids**

Lightsheet video of cleared retinal-brain complex organoids. Video showing 510 z stacks starting at the retinal organoid with typical thick neuroepithelia showing MAP2 positive retinal ganglion cells and axonal projections leading towards cortical organoid with neurons positive for MAP2.

**Supplementary Table 1.** Antibodies used for immunohistochemistry.

| Supplementary Table 1. Antibodies used for immunohistochemistry |  |  |  |
| --- | --- | --- | --- |
| Antigen | Host species | Concentration used | Supplier |
| Calretinin | mouse | 1 in 200 | Merck Millipore (MAB1568) |
| Caspase3 | rabbit | 1 in 200 | Abcam (ab2302) |
| Ctip2 | rat | 1 in 200 | Abcam (Ab18465) |
| FoxG1 | rabbit | 1 in 200 | Abcam (Ab18259) |
| GABA | rabbit | 1 in 200 | Sigma (A2052) |
| GFAP | rat | 1 in 200 | Sigma (345860) |
| Ki67 | rabbit | 1 in 200 | Abcam (ab15580) |
| MAP2 | chicken | 1 in 1000 | Abcam (ab5392) |
| MAP2 | mouse | 1 in 200 | Merck Millipore (MAB3418) |
| NCAD | mouse | 1 in 200 | BD Bioscience (610921) |
| Nestin | mouse | 1 in 200 | Merck Millipore (MAB5326) |
| Pax6 | rabbit | 1 in 200 | Bioscience (901302) |
| PSD95 | goat | 1 in 200 | Abcam (ab12093) |
| Satb2 | rabbit | 1 in 200 | Abcam (Ab34735) |
| SB100 | mouse | 1 in 200 | Sigma (S2532) |
| Sox2 | rabbit | 1 in 200 | Abcam (ab97959) |
| Sox2 | mouse | 1 in 200 | BD (561469) |
| Synaptophysin | mouse | 1 in 200 | Abcam (ab8049) |
| TBR1 | rabbit | 1 in 200 | Abcam (ab31940) |
| Tuj1 | mouse | 1 in 2000 | Thermo Fisher Scientific (MA1118X) |
| Vglut1 | Guinea pig | 1 in 200 | Merck Millipore (AB5905) |

